# Single Platform Solution for Identity and Quantification of Influenza Hemagglutinin (HA) mRNA Constructs and Resulting Expressed Proteins for Application to Influenza mRNA Vaccines

**DOI:** 10.64898/2026.02.05.704054

**Authors:** Rachel Y. Gao, Tianjing Hu, Amber W. Taylor, Keely N. Thomas, Hiromi Muramatsu, Norbert Pardi, Kathy L. Rowlen, Erica D. Dawson

## Abstract

The rapid response to the Coronavirus Disease (COVID-19) pandemic highlighted mRNA as an attractive modality for vaccines and therapeutics to address infectious diseases, cancer, and other diseases. The current analytical methods for identification and quantification of both the mRNA construct as well as the resulting expressed protein(s) relevant to mRNA vaccines are inherently singleplex and usually labor and time intensive, presenting a bottleneck as these vaccines become increasing multivalent. Here, we first present performance data from an approximately 2-hour DNA microarray assay developed specifically to provide identity and quantity measurements for the mRNA constructs present in an influenza virus hemagglutinin (HA)-encoding multivalent mRNA vaccine regardless of the specific strain or codon optimization strategy utilized. The assay functions on both naked and lipid nanoparticle (LNP)-encapsulated mRNAs without the requirement of an upfront mRNA extraction or amplification. Second, we show that a separate ∼2-hour multiplexed immunoassay executed on the same VaxArray Platform can be utilized to measure the expressed proteins produced from these influenza HA mRNAs post-transfection as an *in vitro* potency assay readout. Both assays provide the necessary specificity for each component present in a multivalent mixture and represent new tools in the analytical toolbox for multiplexed mRNA vaccine analytics.

## 1 Introduction

During the COVID-19 pandemic, mRNA technology was highlighted as a promising platform for safe and effective vaccines. Additionally, the rapidity and scalability of mRNA vaccine manufacturing means vaccine manufacturers can more efficiently respond to rapidly mutating pathogens such as SARS-CoV-2 and influenza virus compared to more traditional vaccine technologies such as egg culture, cell culture, and recombinant protein. It took approximately 9 months for the two initial COVID-19 mRNA vaccines, Comirnaty® (Pfizer-BioNTech) and Spikevax® (Moderna),^1-3^ to gain Emergency Use Authorization after the COVID-19 pandemic was declared—an unprecedented speed, particularly in the context of a global pandemic.^2^ Currently, numerous mRNA vaccines are in pre-clinical and clinical development from different manufacturers and academic and government,^4,5^ with over 490 entries under “mRNA” on clinicaltrials.gov for phase 1, 2 or 3 clinical trials.^6^

One advantage of mRNA vaccine technology is that the same 3’- and 5’-untranslated regions (UTRs) and same proprietary lipid nanoparticle (LNP) vaccine delivery system may be applicable to a variety of vaccine targets, with coding sequences easily modified to reflect different antigens or antigenic drift of the included antigens.^2^ The previously licensed bivalent COVID-19 mRNA vaccines showcased that this technology is suitable for the delivery of several unique mRNA constructs, individually LNP-encapsulated, and offered in one final vaccine formulation.^7,8^ Influenza vaccines, traditionally trivalent or quadrivalent, are another target for which the mRNA-based approach is attractive. As of December 15th, 2025, 14 mRNA influenza vaccine candidates are in development: 5 in phase 1, 5 in phase 2, and 4 in phase 3 clinical trials.^9^

Given mRNA vaccines are only recently licensed, it is not surprising that the guidance on methods to measure mRNA critical quality attributes (CQAs), such as identity, concentration, and purity are still evolving.^1,15-17^ As an example, a United States Pharmacopeia (USP) draft guidance document now in its third draft has undergone recent updates as the analytics have matured, and now recommends a variety of methods for determining identity and concentration.^18^ In addition, European Pharmacopoeia (Ph. Eur.) and European Medicines Agency (EMA) have released draft guidelines outlining regulatory expectations for the quality and control of mRNA vaccines.^19,20^ Sequencing can provide full identity information for mRNA constructs, particularly high throughput sequencing which can be applied to multivalent mRNA samples.^18^ However, sequencing suffers from long times to result, requires significant data analysis expertise, and expensive associated equipment and reagents. While reverse transcription-polymerase chain reaction (RT-PCR) can provide an alternative solution for in-process samples with faster turnaround time and less labor cost, upfront sample preparation to eliminate contaminants/inhibitors and extract mRNA from LNPs are still required in encapsulated samples.^16-18^ In terms of quantification, UV-spectroscopy for naked, unencapsulated mRNA and fluorescence-based approaches such as the RiboGreen assay for LNP-encapsulated mRNA are fast and easy. However, these methods measure *total* concentration and cannot discriminate between concentrations of different unique constructs, which is not suitable for multivalent formulations.^17,18^

Methods for measuring expressed protein(s) produced *in vitro* by mRNA constructs post-transfection into an appropriate cell line, typically termed *in vitro* potency or cell-based potency assays, are also critical tools for the development and release of mRNA vaccines to ensure the correct proteins are being made in sufficient and reproducible amounts. The ideal *in vitro* assay will then correlate with the *in vivo* behavior of the mRNA vaccine post-vaccination.^21^ Current USP guidance calls out ELISA as a typical readout for a cell-based potency assay, but other assays can certainly be utilized including Western blot, surface plasmon resonance, and flow cytometry.^18^ Specifically, single radial immunodiffusion (SRID) is the currently accepted potency assay for traditional protein-based influenza vaccines.^18^ For ease of regulatory acceptance, a SRID-based potency assay for an mRNA-based influenza HA vaccine seems ideal, but SRID suffers from insufficient limits of detection to measure small amounts of expressed protein (as opposed to bulk protein-based vaccine antigen). Antibody-based methods for assessing cell-based potency for influenza vaccines should also be stability-indicating for measurement of conformational, functional protein to better ensure correlation to *in vivo* immunogenicity.^18^

Herein, we showcase two different influenza HA assays developed for the VaxArray multiplexed, microarray-based platform for measuring: 1) identity and quantity of the mRNA constructs, and 2) identity and quantity of the expressed proteins. First, we developed a DNA-based microarray assay termed ‘mRNA flu***IQ***’ capable of simultaneous identification and quantification of multiple mRNA constructs in influenza HA-based mRNA vaccine samples in less than 2 hours. The goal was to design a DNA microarray assay that functions in a ‘universal’ manner, regardless of the specific influenza HA protein the mRNA construct for each vaccine component was designed to produce *in vivo*, and regardless of the codon optimization scheme utilized. Herein, we build on our previous work on DNA arrays for mRNA identity and quantity^22,23^ for specific constructs by seeking universal detection using influenza as a case study. Given the ever-evolving nature of the influenza virus, forward-compatibility with future vaccine strain recommendations is desirable to avoid the need to update and re-validate new assays after each semi-annual vaccine strain recommendation. Compatibility with multiple codon optimization strategies increases utility across developers and manufacturers. Our assay demonstrates high specificity, vaccine-relevant detection limits, and accurate and precise quantification for a wide range of LNP-encapsulated, hemagglutinin (HA)-coding mRNA constructs designed from wild-type strains spanning 11 vaccine seasons and representing multiple sources and codon optimization strategies.

We then highlight a second application of the same VaxArray platform in confirming identity and quantifying the expressed proteins post-transfection in HEK-293T cells. An existing VaxArray immunoassay composed of monoclonal antibodies to capture expressed proteins, VaxArray OmniFlu™ HA Assay, is capable of accurate and precise multiplexed quantification of expressed influenza HA proteins in less than 2 hours. In both monovalent and multivalent transfections with Lipofectamine™ or LNP-encapsulated mRNAs, we highlight the utility of this assay as an *in vitro* potency assay readout for use during *in vitro* potency assay optimization, mRNA construct and formulation optimization, and qualitatively for identity confirmation. The availability of a single platform offering assays that are easily customizable and have applications to both the mRNA constructs and expressed proteins will add value for multivalent influenza mRNA vaccine developers and manufacturers at multiple points in their vaccine development programs.

## 2 Materials and Methods

### 2.1 VaxArray mRNA fluIQ Assay-Related Materials and Methods

#### 2.1.1 Oligonucleotide sequence design and mRNA fluIQ microarray printing

Protein sequences for the WHO-recommended influenza vaccine strains from the 2011-2012 through 2023-2024 Northern Hemisphere seasons were downloaded from Global Initiative on Sharing All Influenza Data (GISAID; Munich, Germany) and aligned in BioEdit (v7.2.5, Ibis Therapeutics, Carlsbad, CA). Amino acid regions conserved across all strains of influenza A/H1 (H1), A/H3 (H3), B/Yamagata (B/Y), and B/Victoria (B/V) were identified using Clustal Omega.^24,25^ In addition, conserved amino acid regions shared by both B/Y and B/V strains were also mapped to enable design of captures for pan-influenza B HA detection. Protein sequences were reverse-translated using fundamental codon optimization, with antisense captures designed to bind the RNA sequence. Capture oligos design focused on the conserved amino acid regions to provide specificity to the subtype/lineage specific constructs with further evaluation with OligoAnalyzer™ (IDT; Coralville, IA, USA). Capture oligos were designed to have T_m_ > 41 °C to enable efficient room temperature hybridization and to have low self-dimerization potential (ΔG ≥ 5.5 kcal/mol). Oligos were evaluated *in silico* against off-target mRNA constructs using BLAST (BV-BRC(3.35.5)) to anticipate non-specific interactions.

A total of 126 HA capture oligos were designed and purchased from IDT (Coralville, IA, USA) with a 5’-amino C6 modification and HPLC purification: 30 targeting H1, 32 targeting H3, 18 targeting B/Y, 26 targeting B/V, and 20 targeting both influenza B lineages (“Pan-B”). Capture oligos were printed in 16 replicate microarrays per slide on epoxide-functionalized glass slides at InDevR using a piezoelectric printer. Reactivity and specificity of all 126 oligos were evaluated using a fluorescently conjugated PolyT detection oligo targeting the 3’ polyA tail of hybridized mRNA constructs. Based on these assessments, 32 oligos were selected for the final array with each printed in four replicate spots per microarray.

#### 2.1.2 mRNA Constructs

mRNA constructs utilized are outlined in **Supplementary Table 1** and represent the WHO-recommended strains for all four vaccine components from the Northern Hemisphere 2018-2019 through 2023-2024 seasons. Briefly, to represent the diversity in mRNA constructs utilized by different groups, mRNA constructs were obtained from 3 different sources, with each source utilizing a different set of UTRs and codon optimized with one of three different methods as described below. Full coding region sequences for the constructs summarized in **Supplementary Table 1** are provided in **Supplementary Table 2**.

Source 1 constructs were synthesized at the University of Pennsylvania, including a 101 nucleotide (nt)-long polyA tail and proprietary UTRs, and were prepared as described.^26^ Source 2 constructs from GenScript Biotech (Piscataway, NJ, USA) included a 100 nt polyA tail and proprietary UTRs. Source 3 constructs from TriLink Biotechnologies (San Diego, CA, USA) included a 120 nt polyA tail and proprietary UTRs.

#### 2.1.3 Codon Optimization Strategies

Four different codon optimization strategies were used. Method 1 refers to fundamental codon optimization utilizing the codon recognized by the most abundant (dominant) tRNAs. Method 2 refers to the GenScript Biotech codon optimization approach. Method 3 refers to the GENEWIZ (South Plainfield, NJ, USA) codon optimization approach. Method 4 is a variation on fundamental codon optimization and was only applied to materials from the University of Pennsylvania.

#### 2.1.4 LNP encapsulation of mRNA and encapsulation efficiency

For any mRNA flu***IQ*** assay experiments involving LNP-encapsulated mRNAs, mRNA constructs were encapsulated with Hepato9 mRNA transfection kit (Precision Nanosystems Inc., Vancouver, BC, Canada) using the NanoAssemblr® Spark™ microfluidic mixer (Precision Nanosystems Inc.) following the manufacturer’s recommended protocol.

Encapsulated materials were stored at 2-8 °C prior to analysis, with quantity and encapsulation efficiency determined using the Quanti-it™ RiboGreen RNA Assay Kit (Cat#R11490; Invitrogen, Waltham, MA, USA) per manufacturer’s instructions as described previously.^22^ In short, standard curves and unknown samples were plated in a 96-well format and incubated at 37 °C for 10 min to lyse LNPs prior to RiboGreen addition. Total mRNA was measured in TE buffer containing 1% Triton X-100 (SX100; Sigma-Aldrich), and free mRNA was measured in the absence of detergent. Fluorescence was measured using a FLUOstar OPTIMA plate reader (BMG LABTECH, Ortenberg, Germany) at 480 nm excitation and 520 nm emission, with mRNA quantities calculated from the corresponding standard curves.

#### 2.1.5 mRNA fluIQ Assay Procedure

The VaxArray mRNA flu***IQ*** assay protocol is similar to that previously described.^22,23^ In short, samples were diluted to the target total mRNA concentration by combining 2× Binding Buffer Component A (VX-6320, InDevR, Inc., Boulder, CO, USA), 10× Binding Buffer Component B (VX-6321, InDevR, Inc.), and RNase-free water to achieve 1× final concentrations of components A and B. For LNP-encapsulated samples, lysis was first performed by incubating samples with 1% Tergitol 15-S-9 (15S9, Sigma Aldrich) at 37 °C for 10 min with all subsequent dilutions prepared in mRNA binding buffer supplemented with 1% Tergitol 15-S-9.

Incubations were performed at room temperature in a humidity chamber (VX-6204, InDevR, Inc.). Arrays were pre-washed with 50 µL/array mRNA Wash Buffer A (VX-6324, InDevR, Inc.) for 1 min, the buffer removed, and mRNA samples applied. Slides were incubated for 1 h, washed with mRNA Wash Buffer A for 1 min, and incubated for 15 min with PolyT detection label (VXI-RNA-7601, InDevR, Inc.) diluted in mRNA binding buffer. Slides were subsequently washed sequentially in mRNA Wash Buffer A and mRNA Wash Buffer C (VX-6326, InDevR Inc.), excess liquid was removed by pipette, and slides were spun dry (ArrayDry, (VX-6218, InDevR Inc.) prior to imaging and data analysis using the VaxArray Imaging System (VX-6000, InDevR Inc.) with VaxArray Software version 3.0.16.

#### 2.1.6 Reactivity, Analytical Sensitivity, and Dynamic Range

Capture oligo specificity was screened using monovalent naked mRNA at 20 µg/mL. A capture oligo was considered specific if the signal-to-background (S/B) ratio generated for the on-target mRNA constructs was ≥ 3 and the S/B ratio generated for off-target mRNA constructs was < 3.

To estimate lower and upper limits of quantification (LLOQ and ULOQ, respectively), a 15-point dilution series of monovalent LNP-encapsulated mRNAs starting at 30 μg/mL and a blank were prepared in four replicates from the same stock and processed as described above. For all reactive and specific captures, the average median RFU value at each concentration was plotted, and a moving 4-point linear fit was applied, generating a slope and R^2^ for each 4-point fit. The LLOQ was estimated as the concentration within the lowest 4-point fit for which R^2^ ≥ 0.95 at which the signal was equal to the average blank signal + 5σ (σ is standard deviation of the background signal). If the R^2^ for the 4-point fit at the highest concentrations was not ≥ 0.95, the ULOQ was estimated as the concentration at 90% of the signal of the highest qualifying standard. Otherwise, the ULOQ was stated as being ≥ the highest concentration tested.

#### 2.1.7 Accuracy and Precision

The samples described above to determine LLOQ/ULOQ were also used to evaluate accuracy and precision. One replicate dilution series was used as the standard curve, and the average signal of the other three replicates for a chosen concentration within the linear range was back-calculated against the standard curve, with accuracy determined as % of expected signal and precision expressed as % RSD of the three replicates.

### 2.2 Expressed Protein Assay-Related Materials and Methods

#### 2.2.1 Cell Culture

HEK-293T cells (ATCC CRL-3216; American Type Culture Collection, Manassas, VA, USA) were cultured in Dulbecco’s Modified Eagle Medium (DMEM; 11995-065, Gibco, Waltham, MA, USA) supplemented with 10% fetal bovine serum (FBS; A31604-01, Gibco) and 1× Penicillin-Streptomycin-Glutamine (PSG; 10378-016, Gibco) in adherent cell culture flasks (156499, Nunc™ EasYFlask™). Adherent cells were washed with sterile Dulbecco’s phosphate-buffered saline (10010-023, Gibco), followed by incubation with 1 mL of 0.25% Trypsin-EDTA (25200-056, Gibco) for 1–2 minutes, after which cells were collected and trypsin inactivated with 5 mL of HEK-293T cell medium.

#### 2.2.2 LNP encapsulation of mRNA and encapsulation efficiency

For any expressed protein assay experiments involving LNP-encapsulated mRNAs, constructs were encapsulated with DSPC (P1128, Sigma Aldrich, St. Lousi, MO, USA), cholesterol (C8667, Sigma Aldrich), ALC-0315 (890900O, Avanti Polar Lipids, Alabaster, Alabama, USA), and ALC-0159 (880155P, Avanti Polar Lipids) using a similar ratio to the Pfizer Comirnaty vaccine.^27^ Encapsulated materials were stored at 2–8 °C prior to analysis, and mRNA quantification was performed as described previously (Section 2.1.4).

#### 2.2.3 mRNA Transfection

Cells were seeded in 6- or 12-well plates to achieve 70-90% confluency at the time of transfection. Cells were transfected with Lipofectamine MessengerMAX™ Transfection Reagent (LMRNA015, Invitrogen) or lipid nanoparticles (LNPs) formulated using the NanoAssemblr® Spark™ system (Precision NanoSystems Inc., Vancouver, CA) following the manufacturer’s recommended protocol. Transfections were carried out at 37 °C and 5% CO2 incubator for 24 hours unless otherwise specified. Mock-infected cells were treated identically without the addition of mRNA.

For Lipofectamine transfections, mRNA was diluted in Opti-MEM™ Reduced Serum Medium (31985062, Gibco). Lipofectamine MessengerMAX™ reagent was diluted separately in Opti-MEM™ and incubated for 10 minutes at room temperature. Diluted mRNA and reagent were then combined and incubated for an additional 5 minutes to allow complex formation. Prior to transfection, the growth medium in each well was aspirated and replaced with reduced-serum, antibiotic-free medium (DMEM with 0.1% FBS). The mRNA–lipid complexes were then added to the cells. Plates were returned to a 37 °C, 5% CO2 incubator, and the transfection medium was replaced with fresh complete growth medium (DMEM supplemented with 10% FBS and PSG) after 6 hours.

For LNP transfections, mRNA-LNPs were formulated with lipids as described in section 2.2.2. On the day of transfection, mRNA-LNPs were mixed with DMEM containing 10% FBS and recombinant human ApoE3 (4144-AE, R&D Systems) to achieve a final ApoE3 concentration of 1 µg/mL. This mixture was incubated at 37 °C for 20 minutes to facilitate ApoE3 binding to the LNPs. Prior to transfection, growth medium was aspirated and replaced with antibiotic-free medium (DMEM with 10% FBS). After pre-incubation, the LNP–ApoE3 complexes were gently added to the cells while swirling the plates then returned to the incubator.

#### 2.2.4 Cell Harvest, Sample Clarification, and Protein Measurement

To extract expressed protein, culture medium was aspirated, cells were washed three times with ice-cold PBS, and lysed on ice using Pierce™ IP lysis buffer (87787, Thermo Fischer Scientific, Waltham, MA, USA) supplemented with Halt™ protease inhibitor cocktail (78425, Thermo Fischer Scientific). After a 5-minute incubation on ice, cells were scraped and lysates were transferred to pre-chilled 1.7 mL microcentrifuge tubes.

Lysates were gently mixed to disrupt aggregates and sonicated on ice using a Qsonica Q125-110 sonicator (Newtown, CT, USA) at 20% amplitude for three cycles of a 5-second pulse followed by a 15-second pause. Lysates were centrifuged at 12,000 × g for 3 minutes at 4 °C to remove cellular debris, with clarified supernatant transferred to fresh tubes for analysis.

Total protein concentration was determined using the Pierce™ BCA Protein Assay Kit (Thermo Fisher Scientific). Lysates were diluted 1:5 in PBS, plated in duplicate in 96-well format alongside a BSA standard curve, and incubated with working reagent prior to absorbance measurement at 562 nm using a FLUOstar OPTIMA plate reader (BMG LABTECH). Processed lysates were analyzed immediately or snap frozen in liquid nitrogen and stored at –80 °C. Protein concentrations were calculated from the linear regression of the standard curve.

#### 2.2.5 VaxArray Protein Quantitation

HA protein concentrations in cell lysates were quantified using the VaxArray OmniFlu HA assay by back-calculation against strain-matched recombinant HA standard curves. HEK-293T derived recombinant HA proteins obtained from Sino Biological (Beijing, China) included A/Wisconsin/67/2022 (H1; Cat. 40940-V08H), A/Darwin/6/2021 (H3; 40868-V08H), B/Austria/1359417/2022 (B/V; 40862-V08H), and B/Phuket/3073/2013 (B/Y; 40498-V08H1). Standard curves were prepared by serial dilution of each HA in a matrix-matched diluent containing Protein Blocking Buffer (PBB2.0, VX-6305, InDevR, Inc.), Pierce IP Lysis Buffer, and mock-transfected lysate. HA concentrations for each component in the sample of interest were assessed by comparing fluorescence signals generated against the corresponding standard curves using the VaxArray software

#### 2.2.6 VaxArray OmniFlu HA Immunoassay

The VaxArray OmniFlu HA assay follows a similar overall assay protocol to that previously described.^22,23^ VaxArray OmniFlu HA slides (VXI-7750, InDevR, Inc.) and all reagents were equilibrated to room temperature for 30 minutes. Wash buffers (VX-6303 and VX-6304, InDevR, Inc.) were diluted to 1× in deionized water. Lysates were prepared in a mixture of IP Lysis Buffer and Protein Blocking Buffer 2.0, with the final volume comprising at least 50% PBB2.0. Reference antigens were diluted in a matrix-matched diluent containing PBB2.0, IP Lysis Buffer, and mock-transfected lysate at equivalent protein concentrations.

All incubations were performed on an orbital shaker at 80 rpm. Slides were pre-washed with 50 µL of 1× Wash Buffer 1 for 1 min, followed by application of 50 µL of samples or standards per well and incubation for 2 hours at room temperature. After a 1 min wash with 1× Wash Buffer 1, a mixture of four detection label antibodies (A H1-41, VXI-7641; A H3-38, VXI-7638; B Yam-39, VXI-7639; B Vic-33, VXI-7633; InDevR, Inc.) were diluted to 1× in PBB2.0 (without detergent) and incubated for 30 min. Slides were then washed sequentially with Wash Buffer 1, Wash Buffer 2, 70% ethanol, water, and spin dried (ArrayDry, VX-6218, InDevR, Inc.), then imaged and analyzed using the VaxArray Imaging System (VX-6001, InDevR, Inc.).

## 3 Results and Discussion

### 3.1 mRNA fluIQ Assay Design

To develop an assay enabling broad detection of influenza A H1, H3, and influenza B/Y and B/V constructs despite antigenic drift, conserved HA regions were identified across vaccine-relevant strains spanning multiple Northern Hemisphere seasons (2011-2012 to 2023-2024). Given the removal of B/Y from recent vaccine recommendations,^25^ conserved regions shared between B/Y and B/V were used to design Pan-B captures. From these conserved regions, 126 candidate oligonucleotides were designed and screened for reactivity and specificity across a panel of vaccine-related HA mRNA constructs differing in coding sequences, UTRs, and codon optimization strategies. Based on these assessments, 32 oligonucleotides were selected to ensure robust and specific detection across all targeted subtypes and lineages (**Supplementary Table 3**).

The final microarray layout consists of 16 microarrays per slide (**Figure 1a**), with each of the 32 capture oligonucleotides printed in quadruplicate (**Figure 1b**). The assay detection principle is illustrated in **Figure 1c**. Representative fluorescence images qualitatively highlighting assay specificity and multiplex performance are shown in **Figures 1d–h**.

**Figure 1.**
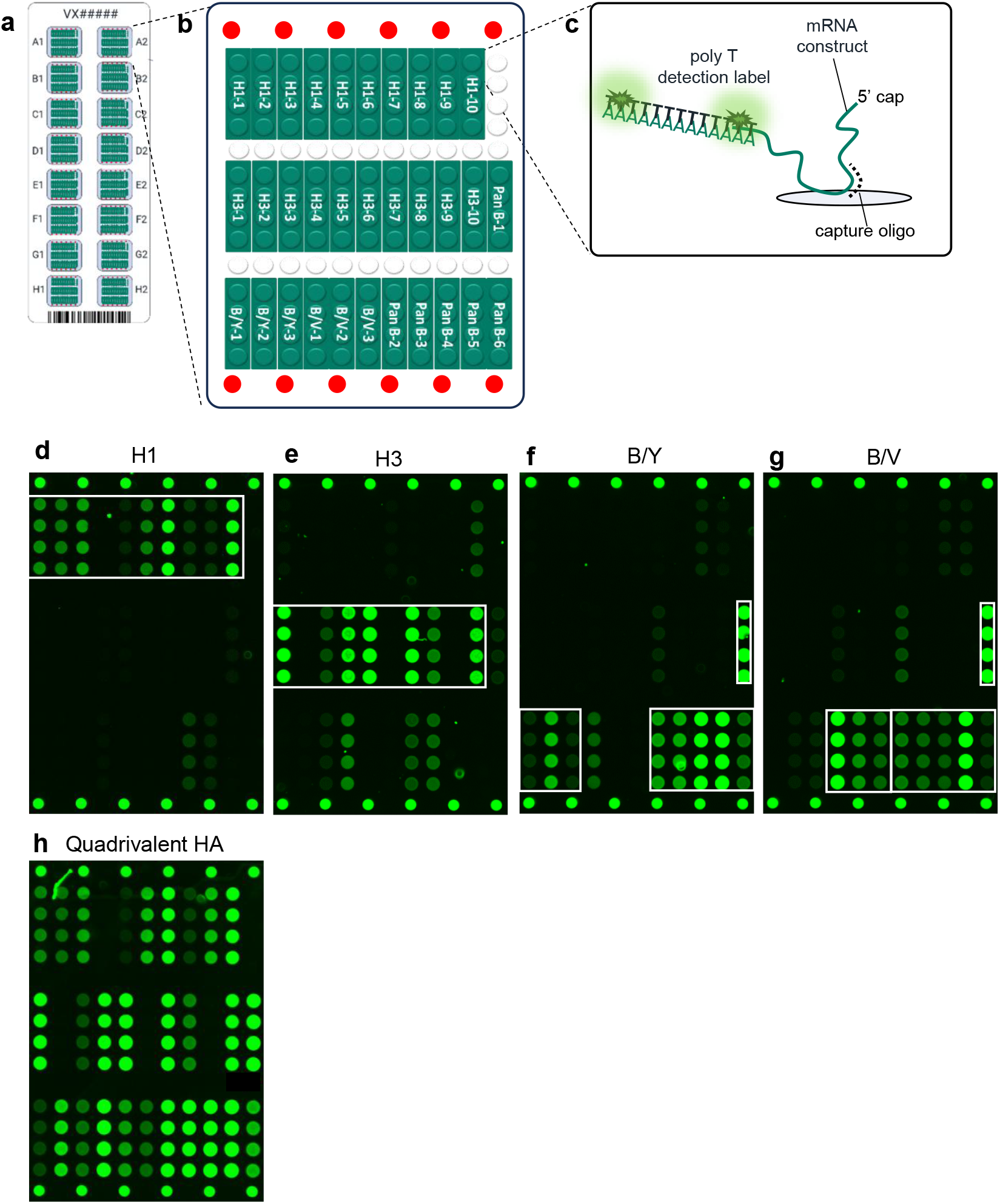
Design of VaxArray flu*IQ* HA Assay. **(a)** Schematic representation of the microarray slide with 16 replicate microarrays, **(b)** individual microarray layout showing 4 replicate spots in vertical columns for each capture oligo as labeled, fiducial markers in red, and white spots for background signal collection, **(c)** assay detection scheme showing capture oligo binding to corresponding mRNA HA-encoding construct and fluorescent polyT detection label. **(d-h)** representative fluorescence images for 10 μg/mL HA-encoding mRNA constructs. **(d)** A/Wisconsin/67/2022 (H1), **(e)** A/Darwin/6/2021 (H3), **(f)** B/Phuket/3073/2013 (B/Y), **(g)** B/Austria/356417/2021 (B/Y), and **(h)** quadrivalent mixture of all 4 mRNA constructs. White boxes highlight target-specific oligos for each monovalent image. Note not all target-specific captures are reactive, and some captures exhibit off-target reactivity.

### 3.2 Assay is reactive and specific to a wide range of targeted mRNA constructs

As shown in **Figure 2**, HA mRNA constructs from Source 3 encoding the Northern Hemisphere 2023–2024 cell-based vaccine strains A/Wisconsin/67/2022 (H1), A/Darwin/6/2021 (H3), B/Phuket/30/2013 (B/Y), and B/Austria/359417/2021 (B/V) were codon-optimized using 3 different methods and evaluated on the microarray at 20 µg/mL. Of the captures designed for each subtype, three H1, two H3, one B/Y, and one B/V capture demonstrated good reactivity, with signal-to-background (S/B) values ranging from 3.1 to 20 across all three intended constructs, and good specificity. Some capture oligos exhibit elevated S/B >2 on off-target constructs, indicating incomplete specificity. The envisioned use case is that an end user assesses monovalent constructs to determine which subset of capture oligos are reactive and specific for each intended target, and that the subset of the overall 32 capture oligos that demonstrate good reactivity and specificity are then down-selected for use in subsequent identity testing and/or quantitative analysis. This overdesign of the microarray to include a wider variety of capture sequences enables the assay to remain reactive and specific over time and across construct design choices.

**Figure 2.**
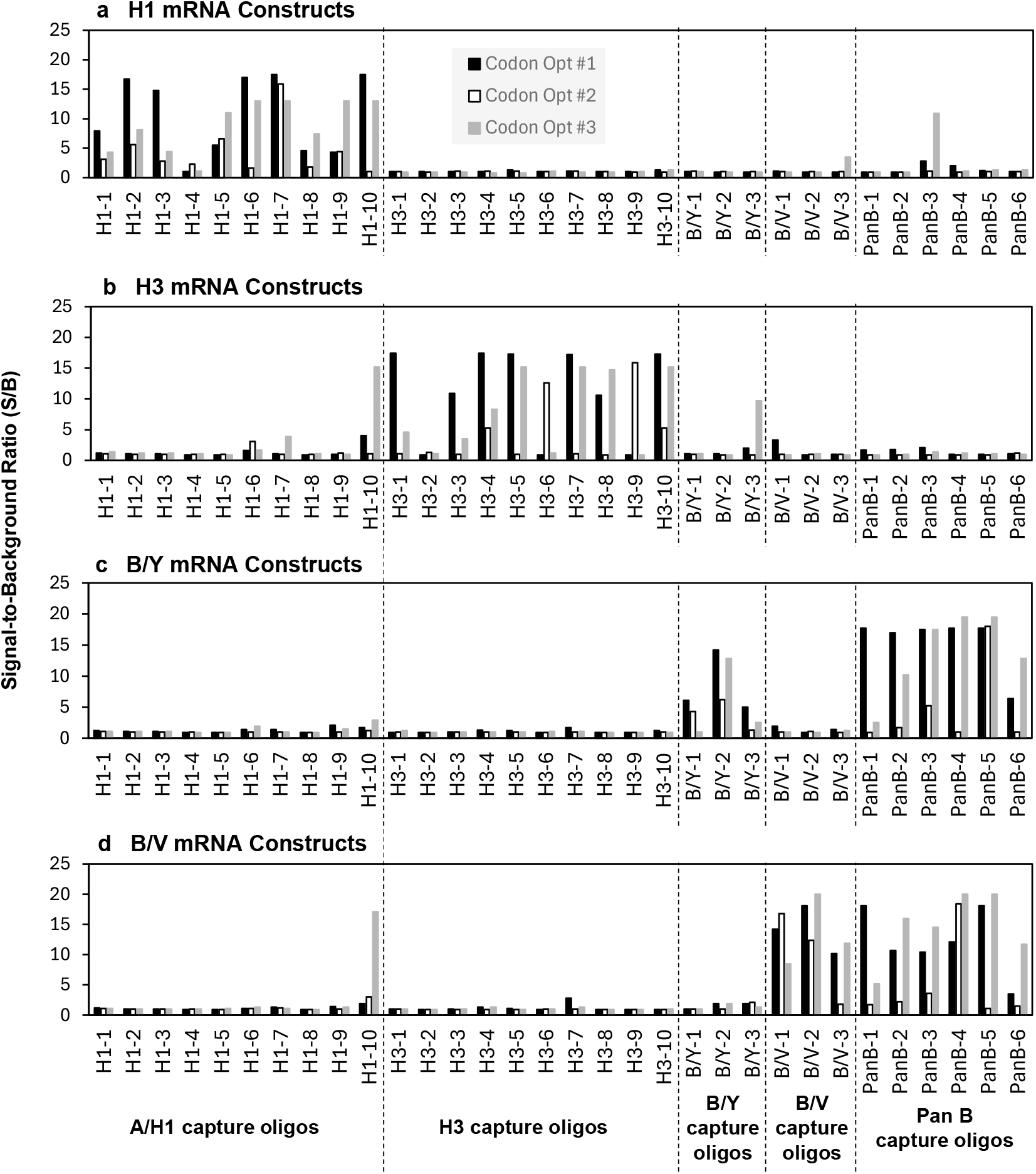
Representative reactivity and specificity of HA-encoding mRNA constructs generated using different codon optimization strategies. Capture oligonucleotides are grouped by target as indicated. Panels a-d highlight constructs encoding the same influenza HA protein but produced using three different codon optimization methods (black, white, and gray series). Panels correspond to **(a)** H1, **(b)** H3, **(c)** B/Y, and **(d)** B/V. All samples were analyzed at 20 µg/mL with a 700 ms exposure time. Signal-to-background (S/B) ratios > 3 on intended captures were considered positive reactivity, while S/B > 3 on off-target captures indicated cross-reactivity. PanB capture oligonucleotides are expected to react with both B/Y and B/V constructs. Data shown are representative of one independent experiment (n = 1). Note these data should be considered qualitative, as high S/B values may be at fluorescence saturation.

To assess the impact of various UTRs on assay performance, HA constructs encoding A/Wisconsin/67/2022 (H1), A/Darwin/6/2021 (H3), B/Phuket/30/2013 (B/Y), and B/Austria/359417/2021 (B/V) with different UTRs were evaluated (**Figure 3)**. In each panel, reactivity varied somewhat for the 3 constructs differing in only the UTR, but a subset of captures was generally reactive to all three, indicating that constructs can be reliably detected regardless of UTR. As shown in **Figure 2**, reasonably good specificity with most S/B ratios <2 for off-target capture oligonucleotides was observed.

**Figure 3.**
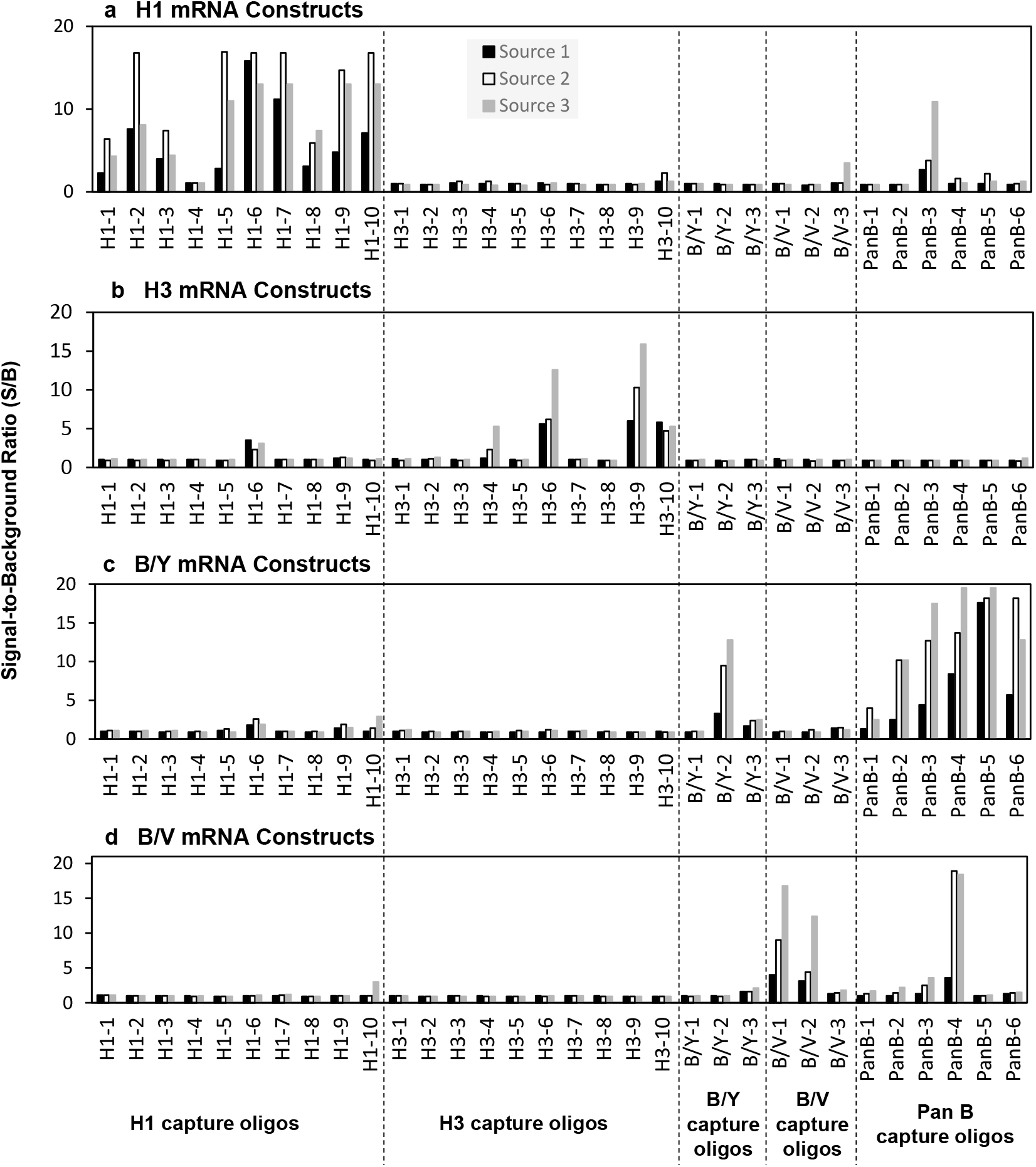
Representative reactivity and specificity of HA-encoding mRNA constructs from different sources using distinct 5′ and 3′ UTRs. Capture oligonucleotides are grouped by target, as indicated at the bottom of each panel. Each plot represents constructs encoding a single influenza A HA subtype or influenza B lineage, with three constructs encoding the same HA protein but derived from different sources and incorporating different UTRs, shown in black, white, and grey. Panels correspond to (a) H1, (b) H3, (c) B/Y, and (d) B/V constructs. All constructs analyzed at 20 µg/mL with 700 ms exposure time. Signal-to-background (S/B) ratios >3 for on-target captures were considered reactive, while S/B ratios >3 for any off-target captures were considered cross-reactive. Pan-B capture oligonucleotides are expected to be reactive with both B/Y and B/V constructs. Data shown are representative of one independent experiment (n = 1). Note these data should be considered qualitative, as high S/B values may be at fluorescence saturation.

Detectable signals were observed for at least one capture across all constructs with differing codon-optimization strategies and UTRs, demonstrating assay tolerance to typical mRNA design modifications. Signal generation relies on polyT binding to the 3’ polyA tail, constructs lacking a sufficiently long polyA tail may show reduced or absent reactivity, and differences in polyA tail properties can contribute to variations in signal intensity, as described previously.^22,23^ Detection of HA constructs representing recent circulating strains further confirms that the selected conserved regions retain relevance despite ongoing antigenic drift. Overall, results shown in **Figures 2 and 3**, and **Supplementary Table 3** demonstrate that the down-selected 32 capture oligos support reliable detection of HA mRNA constructs across circulating influenza A and B lineages, even as seasonal strains evolve, and support the assay’s suitability for broad HA mRNA construct characterization and analytical development by providing a detection framework resilient to antigenic drift.

### 3.3 Comparable Mono- vs. Multivalent Sample Response Indicates No Interference

For all subsequent studies, Method 1 codon-optimized HA mRNA constructs encoding A/Wisconsin/67/2022 (H1), A/Darwin/6/2021 (H3), B/Phuket/3073/2013 (B/Y), and B/Austria/356417/2021 (B/Y) from Source 3 were carried forward, with analysis focused on the subset of capture probes demonstrating strong reactivity and specificity in **Figure 2**.

Although the assay’s high specificity supports identity testing of monovalent constructs, potential interference from other constructs was evaluated to determine the suitability of independent quantification in multivalent mixtures. Each mRNA construct was serially diluted from 20 to 0.09 µg/mL with a buffer-only blank and tested monovalently in four replicates, with signals compared to those obtained at matching concentrations in a quadrivalent mixture. Representative response curves (**Figure 4a–d**) demonstrate similar signals between monovalent and multivalent samples across all concentrations, indicating minimal interference. In addition, **Figure 4** shows linear dose response for all four vaccine components, highlighting suitability for quantification of both mono- and multivalent samples.

**Figure 4.**
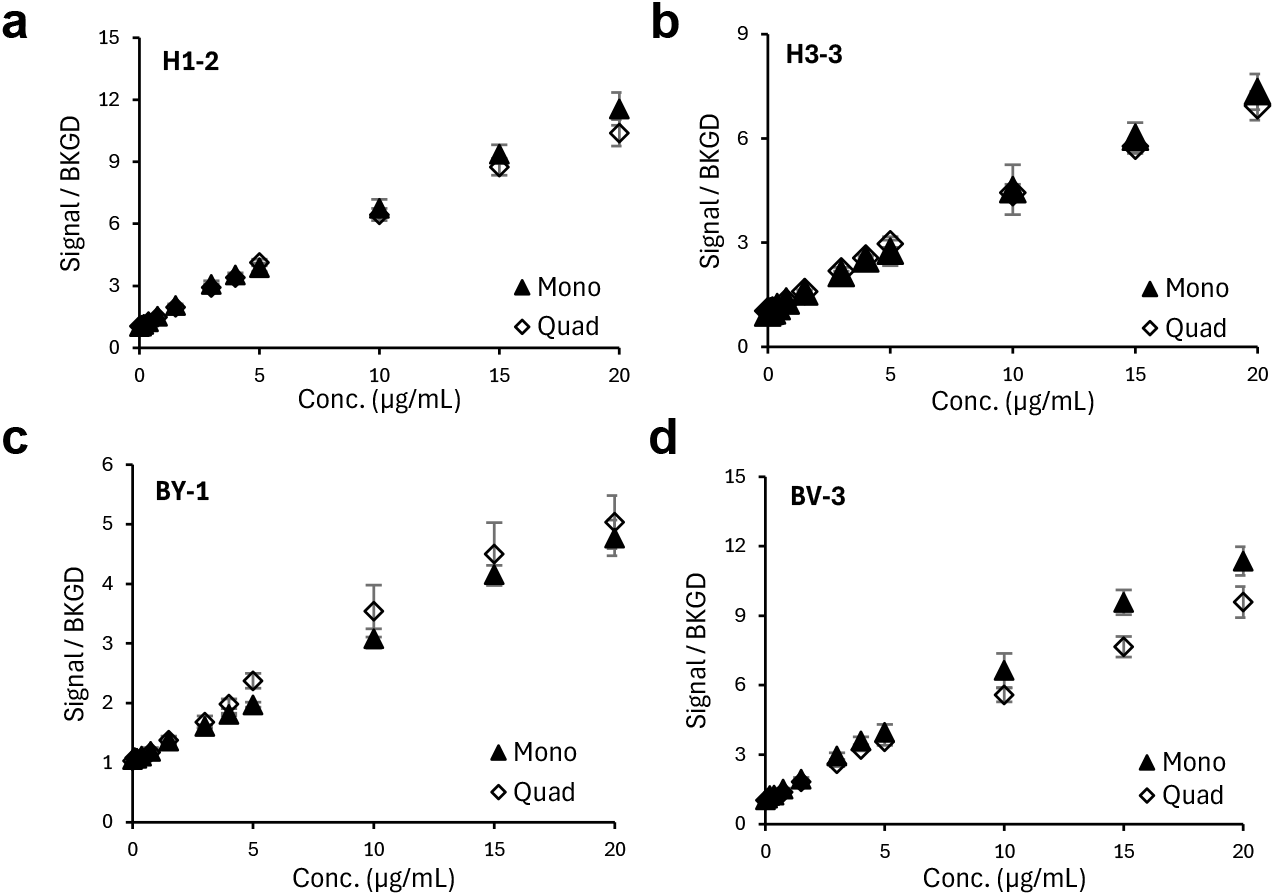
Signal response comparison for monovalent and multivalent naked mRNA materials. Response curves on capture sequences denoted in each panel for Method 1 codon-optimized HA mRNA, both monovalent (▴) and quadrivalent (◊); (a) A/Wisconsin/67/2022 (H1), (b) A/Darwin/6/2021 (H3), (c) B/Phuket/3073/2013 (B/Y), and (d) B/Austria/356417/2021 (B/V). All mRNAs were from source 3. (a-d) Error bars represent one standard deviation (n=4).

### 3.4 VaxArray Assay Performance with LNP-Encapsulated mRNA Constructs

LNP-encapsulated mRNA constructs were evaluated to assess assay performance with vaccine-relevant formulations. Response curves (**Figure 5a–d**) demonstrate that both naked and LNP-encapsulated mRNA generated dose-dependent increases in signal, with encapsulated constructs producing nominally higher signals in some cases at mid-to-high concentrations. At a fixed concentration (4 µg/mL), LNP-encapsulated constructs produced comparable or greater signal relative to quadrivalent naked mRNA across the evaluated captures (**Supplementary Figure 1**), consistent with the monovalent trends. Together, these findings indicate that LNP encapsulation does not impede assay reactivity and that the VaxArray mRNA HA assay can detect and quantify LNP-encapsulated mRNA following a simple lysis step, ideally using a matched LNP standard for quantification.

**Figure 5.**
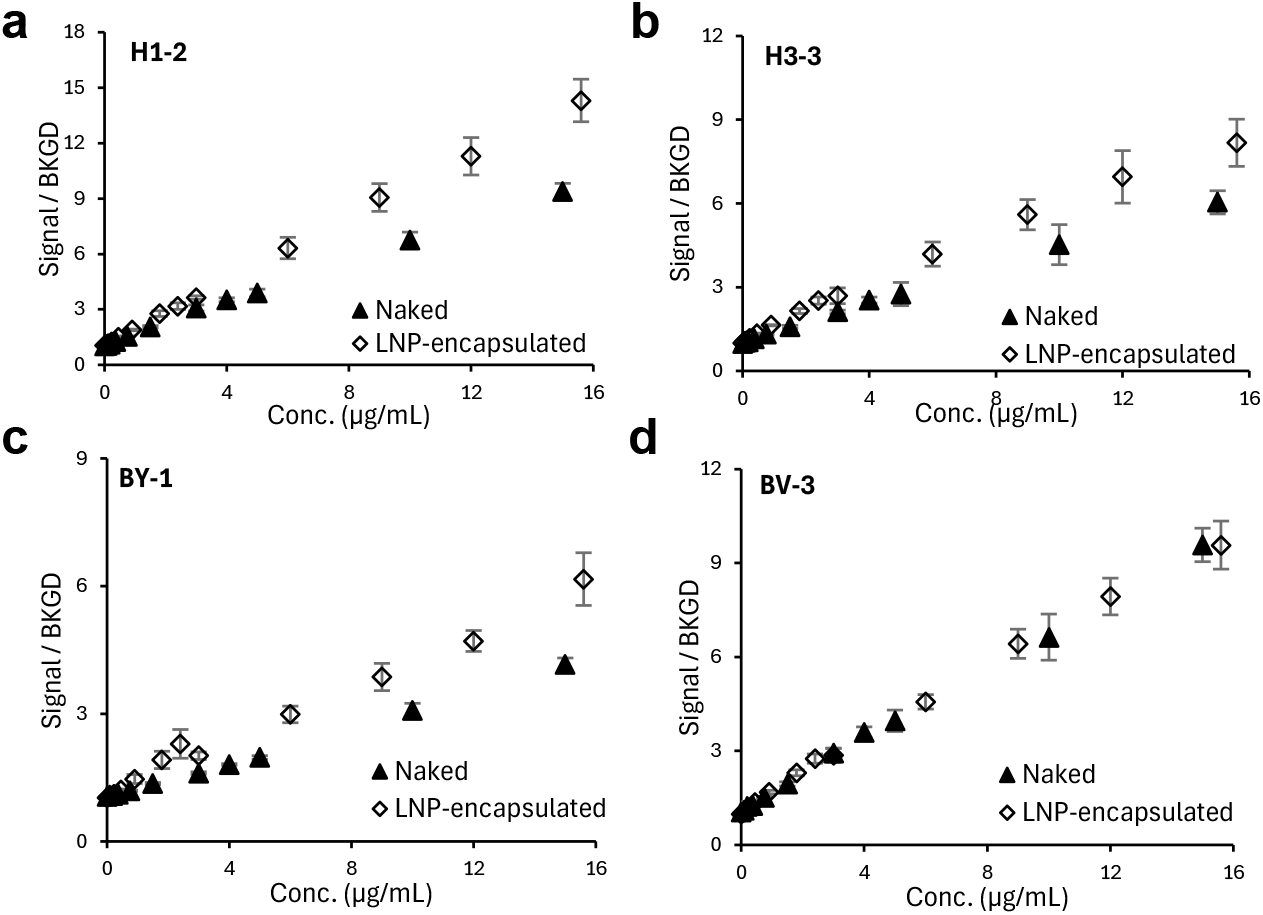
Signal response comparison for monovalent naked and corresponding LNP-encapsulated mRNA materials. Response curves on capture sequences denoted in each panel for Method 1 codon-optimized HA mRNA constructs, both naked (▴) and LNP-encapsulated (◊); (a) A/Wisconsin/67/2022 (H1), (b) A/Darwin/6/2021 (H3), (c) B/Phuket/3073/2013 (B/Y), and (d) B/Austria/356417/2021 (B/V). All mRNAs were from source 3. (a-d) Error bars represent one standard deviation (n=4).

### 3.5 Assay has vaccine-relevant sensitivity and good working range on tested LNP-encapsulated mRNA constructs

The LLOQ and ULOQ were estimated individually for the four HA constructs using LNP-encapsulated mRNA. Fifteen-point monovalent calibration curves ranging from 0.06 to 30 µg/mL were prepared in four replicates and analyzed using captures reactive and specific to each construct (**Supplementary Figure 2**). The LLOQ was estimated as the concentration at which signal equaled the mean blank signal plus 5σ (σ = standard deviation of the background). For all captures evaluated, linear regression of the top four standards yielded R^2^ > 0.95, indicating ULOQ are above the highest tested concentration (>30 µg/mL). Estimated LLOQs, ULOQs, and quantification ranges for each capture are summarized in **Table 1**.

**Table 1.**
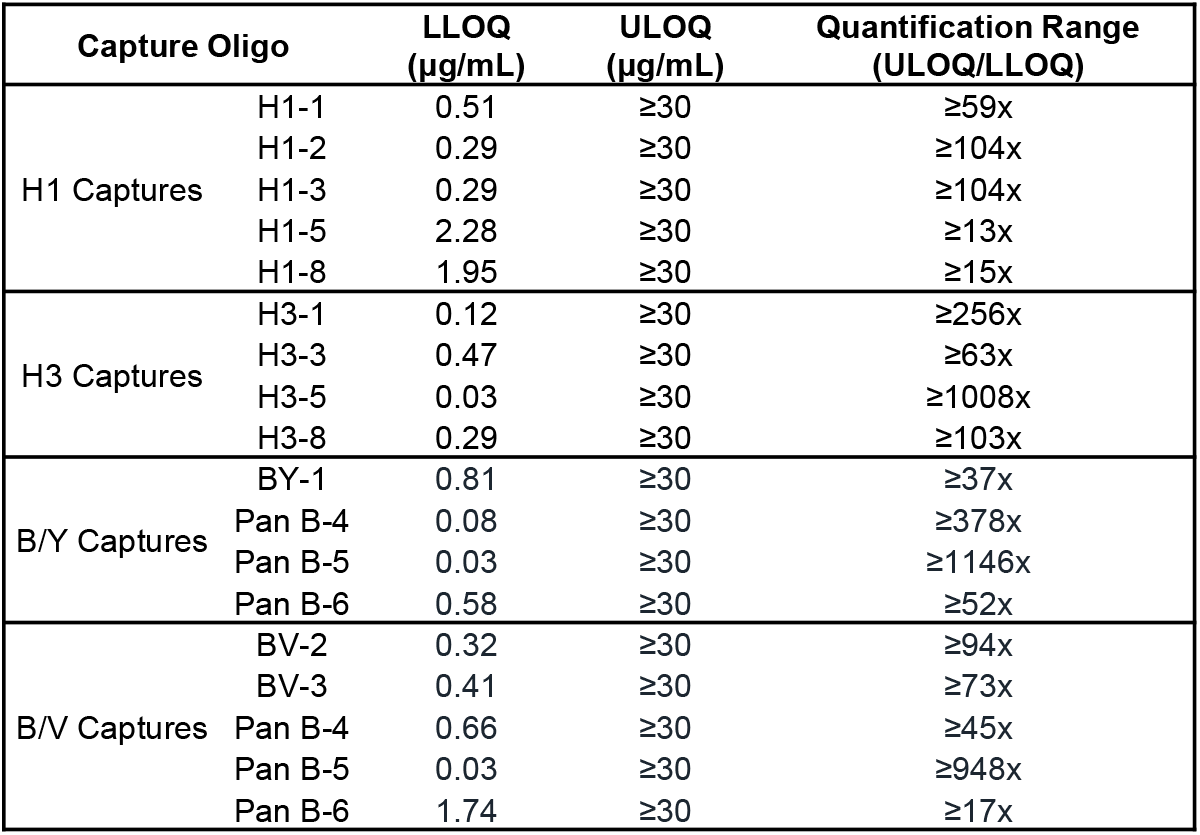
Analytical sensitivity and quantification range of the VaxArray mRNA flu*IQ* assay for LNP-encapsulated HA mRNAs. LLOQ, ULOQ, and quantification ranges for monovalent, LNP-encapsulated A/Wisconsin/67/2022 (H1), A/Darwin/6/2021 (H3), B/Phuket/3073/2013 (B/Y), B/Austria/356417/2021 (B/V) HA mRNAs (Method 1 codon optimization, Source 3) for the representative capture oligos shown. Values were determined from four technical replicates (n = 4).

| Capture Oligo | | LLOQ<br>($\mu\text{g/mL}$) | ULOQ<br>($\mu\text{g/mL}$) | Quantification Range<br>(ULOQ/LLOQ) |
| --- | --- | --- | --- | --- |
| H1 Captures | H1-1 | 0.51 | $\geq 30$ | $\geq 59\text{x}$ |
| | H1-2 | 0.29 | $\geq 30$ | $\geq 104\text{x}$ |
| | H1-3 | 0.29 | $\geq 30$ | $\geq 104\text{x}$ |
| | H1-5 | 2.28 | $\geq 30$ | $\geq 13\text{x}$ |
| | H1-8 | 1.95 | $\geq 30$ | $\geq 15\text{x}$ |
| H3 Captures | H3-1 | 0.12 | $\geq 30$ | $\geq 256\text{x}$ |
| | H3-3 | 0.47 | $\geq 30$ | $\geq 63\text{x}$ |
| | H3-5 | 0.03 | $\geq 30$ | $\geq 1008\text{x}$ |
| | H3-8 | 0.29 | $\geq 30$ | $\geq 103\text{x}$ |
| B/Y Captures | BY-1 | 0.81 | $\geq 30$ | $\geq 37\text{x}$ |
| | Pan B-4 | 0.08 | $\geq 30$ | $\geq 378\text{x}$ |
| | Pan B-5 | 0.03 | $\geq 30$ | $\geq 1146\text{x}$ |
| | Pan B-6 | 0.58 | $\geq 30$ | $\geq 52\text{x}$ |
| B/V Captures | BV-2 | 0.32 | $\geq 30$ | $\geq 94\text{x}$ |
| | BV-3 | 0.41 | $\geq 30$ | $\geq 73\text{x}$ |
| | Pan B-4 | 0.66 | $\geq 30$ | $\geq 45\text{x}$ |
| | Pan B-5 | 0.03 | $\geq 30$ | $\geq 948\text{x}$ |
| | Pan B-6 | 1.74 | $\geq 30$ | $\geq 17\text{x}$ |

The LLOQs ranged from 0.03 to 2.28 μg/mL, with an average of 0.61 μg/mL. ULOQs exceeding 30 µg/mL for all captures, and quantification ranges spanned from ≥13-fold to ≥1146-fold, with the best-performing captures achieving ≥104-fold for H1, ≥1008-fold for H3, ≥1146-fold for B/Y (Pan B-5), and ≥948-fold for B/V (Pan B-5). Given that the mRNA concentration in the currently marketed Pfizer Comirnaty vaccine is 100 µg/mL,^7^ the assay’s quantification range is vaccine-relevant. The assay provides a robust, construct-independent quantification range suitable for mRNA vaccine formulations.

### 3.6 Assay shows good accuracy and precision on tested LNP-encapsulated mRNA constructs

The same four 15-point dilution curves of monovalent LNP-encapsulated mRNA described above were used to assess assay precision and accuracy on relevant (reactive and specific) capture oligos. For each construct, one replicate of the 15-point curve was used as the calibration curve, and three other replicates in the experiment were quantified against it to determine within-experiment precision and accuracy. For each capture, concentrations near the estimated LLOQ (low), the midpoint (medium), and the upper range (high) were selected for back-calculation.

As shown in **Supplementary Table 4**, all selected captures chosen for their reactivity and specificity exhibited high precision, with relative standard deviation below 8.3%, and strong accuracy, with an average of 99.1±9.9% of expected concentration. Only 1 out of the 18 total selected captures produced an accuracy value outside of an acceptable ±20% of expected criterion, and only at a single tested concentration. Overall, these results indicate that the assay demonstrates acceptable precision and accuracy across relevant concentration ranges, with multiple capture options available to support reliable subtype-specific quantification.

### 3.7 VaxArray OmniFlu HA Immunoassay

The VaxArray OmniFlu HA assay is an off-the-shelf, multiplexed microarray-based immunoassay developed by InDevR, Inc. for quantitative measurement of influenza HA antigens and was used as recommended in the instructions for use. The microarray comprises subtype- and lineage-specific capture antibodies printed in a defined layout on functionalized glass slides, enabling simultaneous detection of multiple influenza A subtypes and influenza B lineages, with replicate spots and arrays to support assay robustness. A schematic of the array layout and assay principle is shown in **Figure 6a–c**. In the assay, HA antigens bind to immobilized capture antibodies and are detected using a fluorescently labeled detection antibody, producing signals proportional to concentration of captured HA. Fluorescence is quantified using the VaxArray Imaging System and associated software. We note that other VaxArray influenza HA assays have been previously described and validated for influenza vaccine and antigen characterization,^28-30^ and the OmniFlu HA Assay follows a similar standardized VaxArray workflow to published assays.

**Figure 6.**
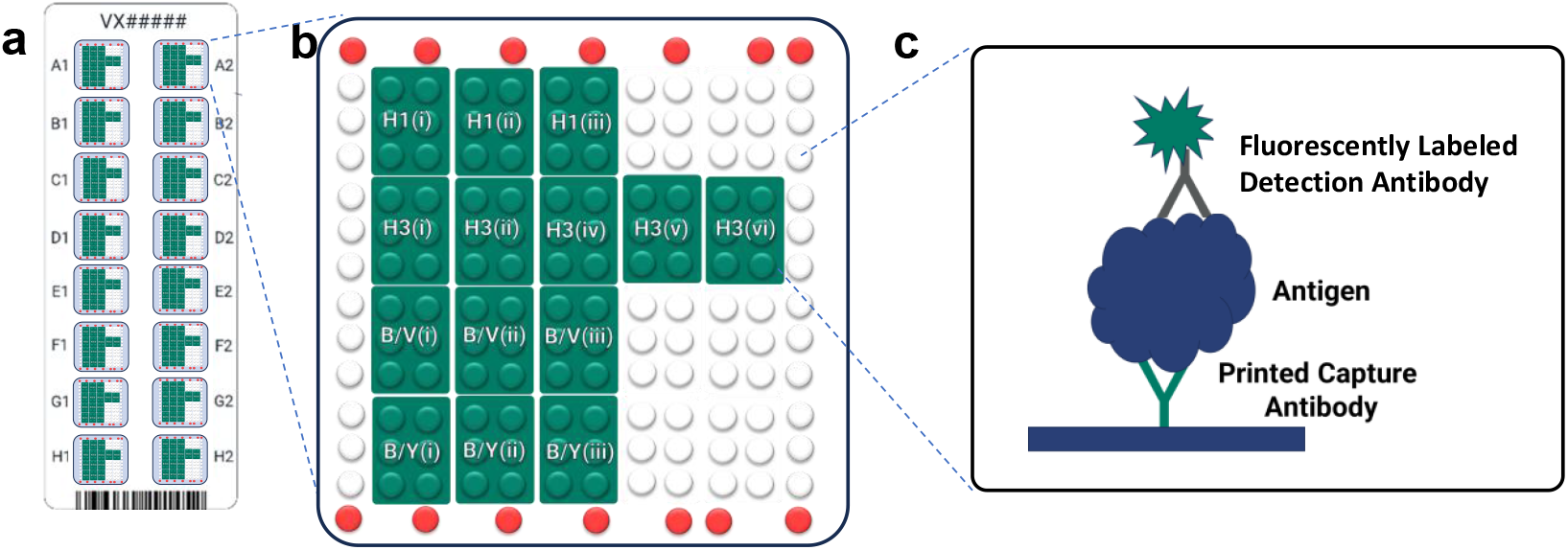
Design of VaxArray OmniFlu Assay. (a) Schematic representation of the VaxArray OmniFlu microarray slide containing 16 replicate microarrays. (b) Layout of an individual microarray showing capture antibody spots printed in green (six replicates per antibody), fiducial markers in red, and background reference spots in white. **(c)** General assay detection scheme in which subtype-specific capture antibodies bind their corresponding HA antigens, followed by detection with a fluorescently labeled secondary antibody.

### 3.8 Detection of Lipofectamine Transfection-Derived HA Protein Across Influenza Subtypes

Quadrivalent mRNA samples (Method 1 codon-optimized mRNA constructs from Source 3, encoding HA from A/Wisconsin/67/2022 (H1), A/Darwin/6/2021 (H3), B/Phuket/3073/2013 (B/Y), and B/Austria/356417/2021 (B/Y)) were transfected in duplicate using Lipofectamine, with each HA mRNA component present at amounts between 0.001 µg to 5 µg per construct (0.004–20 µg total mRNA). Cell lysates were normalized to a fixed total protein concentration of 0.5 mg/mL prior to analysis using the VaxArray OmniFlu HA assay, with three technical replicates analyzed for each biological replicate.

HA concentrations were quantified using the VaxArray OmniFlu HA assay and back-calculated against recombinant HA protein standards included on each assay run. Parallelism between recombinant HA standards and lysate dilution series was evaluated to confirm the appropriateness of the standards and to guide selection of the capture used for data reporting. As shown in **Figure 7a-d**, measured HA concentrations for all four components increased in a dose-dependent manner with increasing mRNA input. In addition, the two biological replicates (separate unique identical transfections) show similar results in terms of protein generated, with good reproducibility across the three technical replicates run at each concentration, demonstrating reproducibility and utility of the assay for measuring expressed protein during optimization of mRNA transfection protocols.

**Figure 7.**
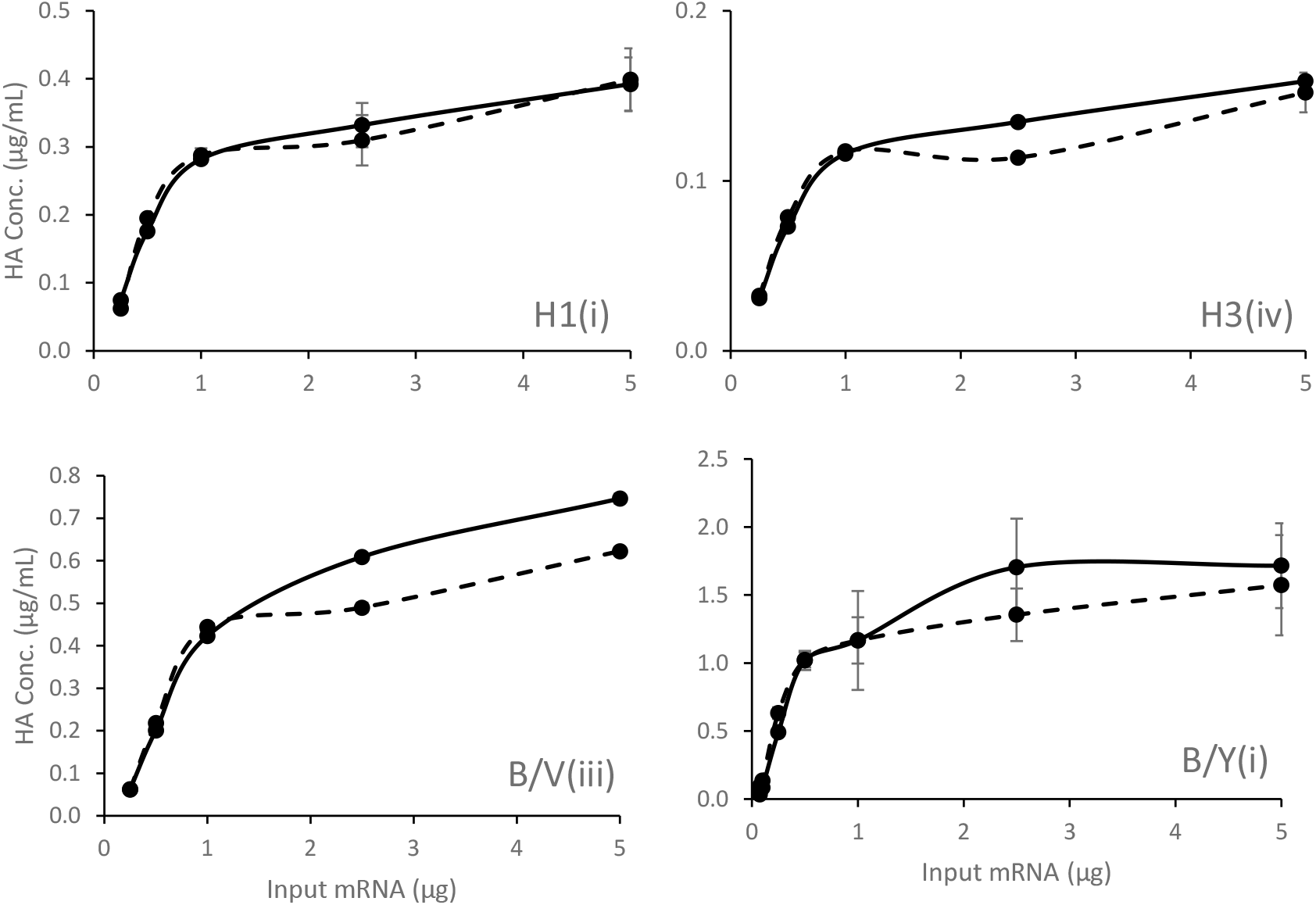
HA protein expression following Lipofectamine transfection of quadrivalent mRNA as a function of input mRNA dose. Quadrivalent mRNA samples encoding A/Wisconsin/67/2022 (H1; 200 ms), A/Darwin/6/2021 (H3; 700 ms), B/Austria/356417/2021 (B/V; 100 ms), and B/Phuket/3073/2013 (B/Y; 400 ms) were transfected across a range of input doses. Expressed HA assessed using the VaxArray OmniFlu HA assay with concentrations determined via back-calculation against recombinant HA protein standard curves. Solid and dashed lines represent two biological replicates; error bars indicate the mean ± standard deviation of technical triplicates (n=3).

### 3.9 Quantitative Detection of HA Protein Expressed from LNP-Formulated mRNA

The same Method 1 codon-optimized A/Wisconsin/67/2022 (H1) mRNA construct from Source 3 was encapsulated into lipid nanoparticles using used in Pfizer’s Comirnaty vaccine formulation.^27^ This monovalent LNP-encapsulated H1 mRNA was transfected across a range of input mRNA amounts. Following transfection, cell lysates were normalized to a fixed total protein concentration of 0.5 mg/mL and analyzed using the VaxArray OmniFlu HA assay, with three technical replicates analyzed from a single biological replicate.

Similar to the Lipofectamine-transfected materials shown in **Figure 7, Figure 8** shows measured H1 HA concentrations increased with increasing input mRNA, demonstrating the dose-dependence of expressed protein as a function of LNP-formulated mRNA. HA protein levels were quantified by back-calculation against recombinant H1 HA protein standards included on each assay run. In addition, the three technical replicates show good reproducibility, with average % RSD of 16.5%. These data indicate the assay can provide a reproducible measurement of HA protein expression after transfection of LNP-encapsulated mRNA formulated using vaccine-relevant lipid compositions for suitability as a readout for expressed protein(s) production post-transfection.

**Figure 8.**
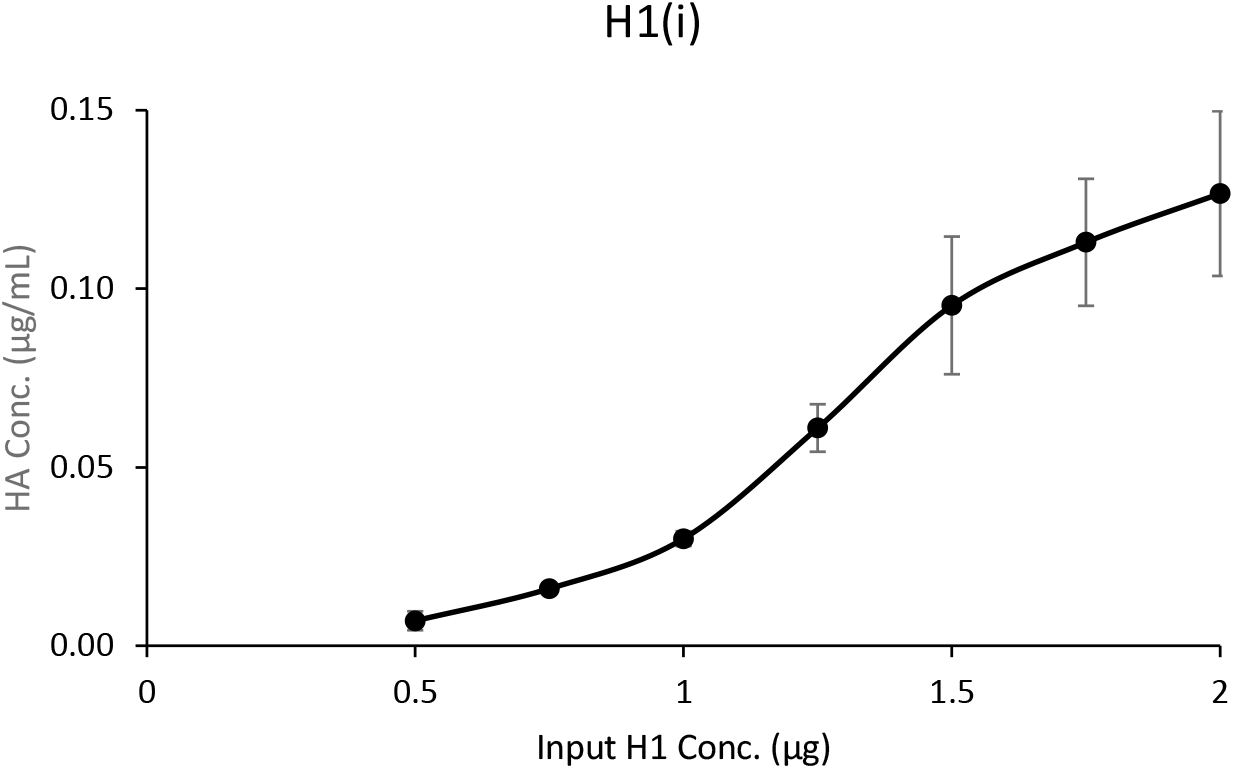
HA protein expression following transfection of monovalent mRNA for H1 HA. Monovalent A/Wisconsin/67/2022 (H1)-coding mRNA was transfected across a range of input doses and resulting HA protein levels in normalized cell lysates were quantified using the VaxArray OmniFlu HA assay. Concentration of expressed HA increased with increasing mRNA input dose. Error bars indicate the mean ± stdev of technical triplicates (n=3).

## 4 Conclusion

The rapidly expanding mRNA therapeutic landscape, including vaccines and other treatments targeting infectious diseases, cancers, and rare diseases presents increasing analytical challenges, particularly for multivalent mRNA drug products. Platform solutions capable of multiplexed analysis that can be utilized at multiple stages in the vaccine development process will serve to help further streamline mRNA vaccine analytics.

In this study, we first demonstrated a multiplexed VaxArray mRNA flu***IQ*** assay capable of sensitive and specific detection of four influenza HA-encoding mRNA constructs with known sequences using fully complementary capture oligonucleotides. The assay enables identity testing as well as accurate and precise quantification of both naked and LNP-encapsulated mRNA in monovalent and multivalent formulations without the need for an additional mRNA extraction step. While RT-PCR methods can support identification and quantification of multivalent mRNA drug substances during manufacturing, they typically require time-consuming mRNA extraction steps for LNP-encapsulated drug products.^16-18^ The constellation of capture sequences on the mRNA flu***IQ*** assay can be down-selected to a set that provides specificity and good reactivity for the constructs of interest in a given vaccine season, with the assay showing specificity and reactivity for subsets of constructs over time indicating some robustness against antigenic drift.

Secondly, we showed that the separate OmniFlu HA assay developed for the same VaxArray platform can be used to provide quantitative measurements of influenza HA proteins expressed following mRNA transfection, including transfection with mRNA-LNPs using a vaccine-relevant lipid formulation. Using influenza HA-based mRNA vaccines as a case study, the data presented herein support the use of VaxArray as a readout for *in vitro* potency as an alternative to other methods such as flow cytometry.

Importantly, while influenza HA mRNA vaccines have been presented as a case study, we note that VaxArray assays addressing other multivalent mRNA vaccines can be easily designed and validated. These capabilities align with emerging needs for standardized analytical methods across mRNA drug substance and drug product characterization workflows in the face of evolving regulatory expectations for mRNA products, and we hope these assays will find increasing utility.

## Data Availability Statement

All relevant data from this study are available from the corresponding author upon request.

## Acknowledgements

The authors have no acknowledgements to declare.

## Author Contributions

RG: Supervision; methodology; investigation; validation; formal analysis; visualization; writing-original draft preparation; writing-review and editing

TH: Supervision; methodology; investigation; validation; formal analysis; visualization; writing-original draft preparation; writing-review and editing

AT: Supervision; methodology; formal analysis; visualization; writing-review and editing

KT: Methodology; investigation; validation; formal analysis; writing-review and editing

HM: Writing-review and editing; resources

NP: Writing-review and editing; resources

KR: Conceptualization; project administration; resources; writing-review and editing

ED: Conceptualization; project administration; resources; supervision; visualization; methodology; funding acquisition; writing-original draft; writing-review and editing

All authors have read and agreed to the published version of the manuscript.

