## Supplemental Figures for "Single Platform Solution for Identity and Quantification of Influenza Hemagglutinin (HA) mRNA Constructs and Resulting Expressed Proteins for Application to Influenza mRNA Vaccines"

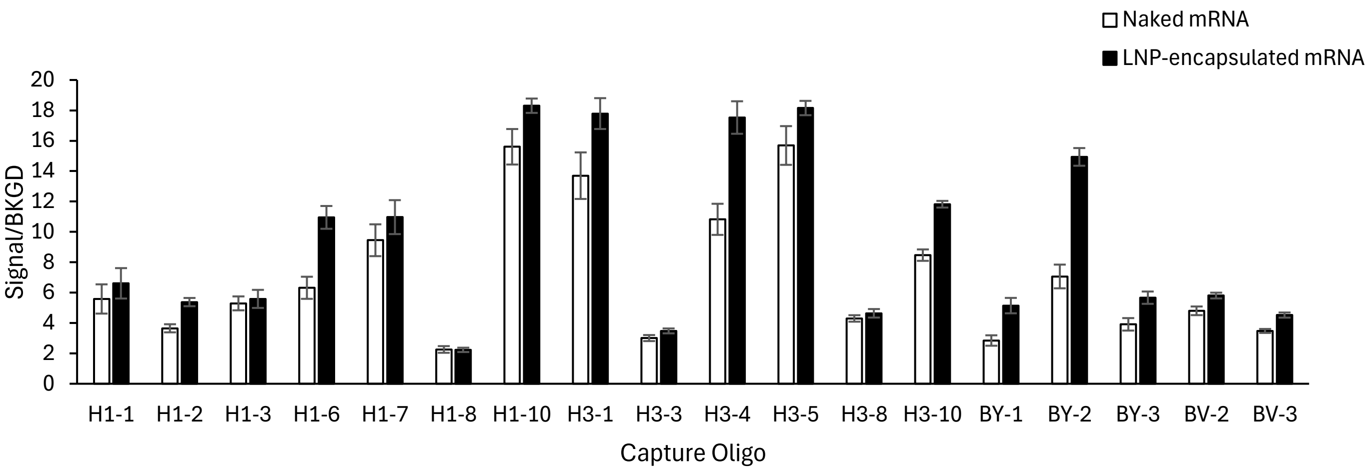


**Supplementary Figure 1. Signal response comparison for naked and LNP-encapsulated mRNA materials in a multivalent mixture.** Method 1 codon-optimized from Source 3 HA mRNA constructs encoding A/Wisconsin/67/2022 (H1), A/Darwin/6/2021 (H3), B/Phuket/3073/2013 (B/Y), and B/Austria/356417/2021 (B/V) (Source 3) tested at 4 µg/mL and 700 ms exposure. Signal response is reported as signal-to-background ratio (Signal/BKGD). Error bars indicate one standard deviation (n = 4).

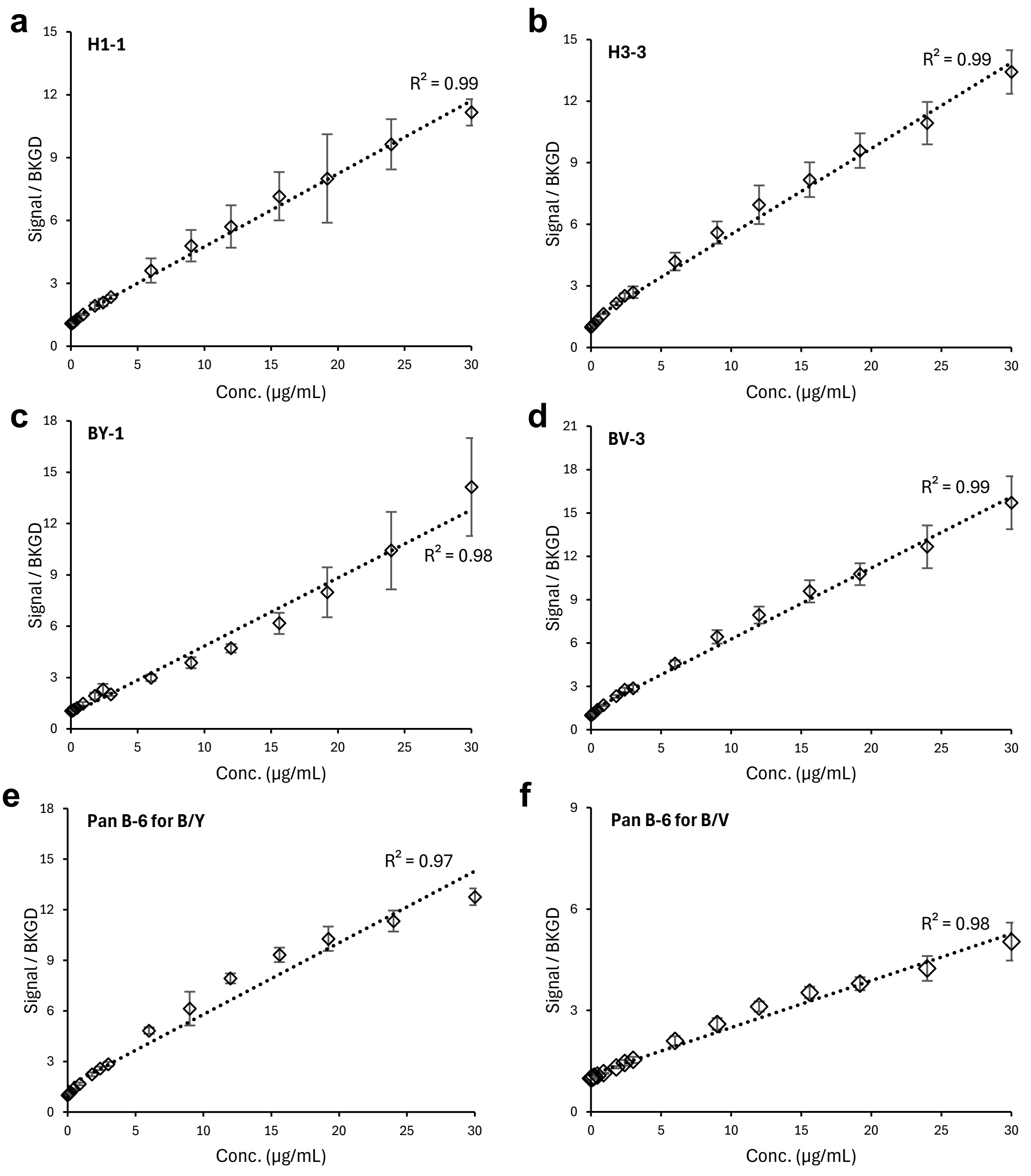


**Supplementary Figure 2. Response curves for LNP-encapsulated HA mRNA constructs.** (a-f) 16-pt serial dilution curves and associated linear regressions for monovalent LNP-encapsulated HA mRNAs (◊) encoding A/Wisconsin/67/2022 (H1), A/Darwin/6/2021 (H3), B/Phuket/3073/2013 (B/Y), and B/Austria/356417/2021 (B/V) (Method 1 codon optimized, Source 3) on the capture oligos shown. Error bars indicate one standard deviation (n=4).
