## Supplemental Table 1 for "Single Platform Solution for Identity and Quantification of Influenza Hemagglutinin (HA) mRNA Constructs and Resulting Expressed Proteins for Application to Influenza mRNA Vaccines"

| Type/ Subtype or Lineage | Strain | Strain included in vaccine season(s): | | | | | | | | | | | | Codon Optimization Method | Source |
|---|---|---|---|---|---|---|---|---|---|---|---|---|---|---|---|
| | | 2018 SH | 2018-2019 NH | 2019 SH | 2019-2020 NH | 2020 SH | 2020-2021 NH | 2021 SH | 2021-2022 NH | 2022 SH | 2022-2023 NH | 2023 SH | 2023-2024 NH | | |
| H1 | A/Wisconsin/67/2022 | | | | | | | | | | | | x | 1 | Source 3 |
| | A/Wisconsin/67/2022 | | | | | | | | | | | | x | 2 | Source 3 |
| | A/Wisconsin/67/2022 | | | | | | | | | | | | x | 3 | Source 3 |
| | A/Wisconsin/67/2022 | | | | | | | | | | | | x | 3 | Source 1 |
| | A/Wisconsin/67/2022 | | | | | | | | | | | | x | 3 | Source 2 |
| | A/Wisconsin/67/2022 | | | | | | | | | | | | x | 2 | Source 2 |
| | A/Sydney/5/2021 | | | | | | | | | | | x | | 2 | Source 2 |
| | A/Wisconsin/588/2019 | | | | | | | x | x | x | x | | | 2 | Source 2 |
| H3 | A/Darwin/6/2021 | | | | | | | | | x | x | x | x | 1 | Source 3 |
| | A/Darwin/6/2021 | | | | | | | | | x | x | x | x | 2 | Source 3 |
| | A/Darwin/6/2021 | | | | | | | | | x | x | x | x | 3 | Source 3 |
| | A/Darwin/6/2021 | | | | | | | | | x | x | x | x | 3 | Source 1 |
| | A/Darwin/6/2021 | | | | | | | | | x | x | x | x | 2 | Source 1 |
| | A/Darwin/6/2021 | | | | | | | | | x | x | x | x | 2 | Source 2 |
| | A/Cambodia/e826360/2020 | | | | | | | | x | | | | | 2 | Source 2 |
| | A/HongKong/45/2019 | | | | | | x | x | | | | | | 2 | Source 2 |
| B/Y | B/Phuket/3073/2013 | x | x | x | x | x | x | x | x | x | x | x | x | 1 | Source 3 |
| | B/Phuket/3073/2013 | x | x | x | x | x | x | x | x | x | x | x | x | 2 | Source 3 |
| | B/Phuket/3073/2013 | x | x | x | x | x | x | x | x | x | x | x | x | 3 | Source 3 |
| | B/Phuket/3073/2013 | x | x | x | x | x | x | x | x | x | x | x | x | 3 | Source 1 |
| | B/Phuket/3073/2013 | x | x | x | x | x | x | x | x | x | x | x | x | 3 | Source 2 |
| B/V | B/Austria/359417/2021 | | | | | | | | | x | x | x | x | 1 | Source 3 |
| | B/Austria/359417/2021 | | | | | | | | | x | x | x | x | 2 | Source 3 |
| | B/Austria/359417/2021 | | | | | | | | | x | x | x | x | 3 | Source 3 |
| | B/Austria/359417/2021 | | | | | | | | | x | x | x | x | 3 | Source 1 |
| | B/Austria/359417/2021 | | | | | | | | | x | x | x | x | 2 | Source 1 |
| | B/Austria/359417/2021 | | | | | | | | | x | x | x | x | 2 | Source 2 |
| | B/Washington/02/2019 | | | | | x | x | x | x | | | | | 2 | Source 2 |

**Supplementary Table 1. Summary of mRNA constructs used in mRNA flu*IQ* assay testing, including strain, vaccine season, and codon optimization method, and construct source.** NH=Northern Hemisphere, SH=Southern Hemisphere. Full coding sequences can be found in **Supplementary Table 2.**
