## Supplemental Table 2 for "Single Platform Solution for Identity and Quantification of Influenza Hemagglutinin (HA) mRNA Constructs and Resulting Expressed Proteins for Application to Influenza mRNA Vaccines"

**Supplementary Table 2: Coding region nucleotide sequences for all influenza HA mRNA constructs evaluated across subtypes and lineages.**

**Type/Subtype:** H1

**Strain:** A/Wisconsin/67/2022
**Codon Optimization Scheme:** 1
**Source:** 3
ATGAAGGCCATCCTGGTGGTGATGCTGTACACCTTCACCACCGCCAACGCCGACACCCTGTGCATCG
GCTACCACGCCAACAACAGCACCGACACCGTGGACACCGTGCTGGAGAAGAACGTGACCGTGACC
CACAGCGTGAACCTGCTGGAGGACAAGCACAACGGCAAGCTGTGCAAGCTGCGCGGCGTGGCCCC
CCTGCACCTGGGCCAGTGCAACATCGCCGGCTGGATCCTGGGCAACCCCGAGTGCGAGAGCCTGA
GCACCGCCCGCAGCTGGAGCTACATCGTGGAGACCAGCAACAGCGACAACGGCACCTGCTACCCC
GGCGACTTCATCAACTACGAGGAGCTGCGCGAGCAGCTGAGCAGCGTGAGCAGCTTCGAGCGCTTC
GAGATCTTCCCCAAGACCAGCAGCTGGCCCAACCACGACAGCGACAACGGCGTGACCGCCGCCTG
CAGCCACGCCGGCGCCCGCAGCTTCTACAAGAACCTGATCTGGCTGGTGAAGAAGGGCAAGAGCTA
CCCCAAGATCAACCAGACCTACATCAACGACAAGGGCAAGGAGGTGCTGGTGCTGTGGGGCATCCA
CCACCCCCCCACCATCACCGACCAGGAGAGCCTGTACCAGAACGCCGACGCCTACGTGTTCGTGG
GCACCAGCCGCTACAGCAAGAAGTTCAAGCCCGAGATCGCCACCCGCCCCAAGGTGCGCGACCAG
GCCGGCCGCATGAACTACTACTGGACCCTGGTGGAGCCCGGCGACAAGATCACCTTCGAGGCCAC
CGGCAACCTGGTGGCCCCCCGCTACGCCTTCACCATGGAGAAGGAGGCCGGCAGCGGCATCATCA
TCAGCGACACCCCCGTGCACGACTGCAACGCCACCTGCCAGACCCCCGAGGGCGCCATCAACAC
CAGCCTGCCCTTCCAGAACGTGCACCCCATCACCATCGGCAAGTGCCCCAAGTACGTGCGCAGCAC
CAAGCTGCGCCTGGCCACCGGCCTGCGCAACGTGCCCAGCATCCAGAGCCGCGGCCTGTTCGGC
GCCATCGCCGGCTTCATCGAGGGCGGCTGGACCGGCATGGTGGACGGCTGGTACGGCTACCACCA
CCAGAACGACCAGGGCAGCGGCTACGCCGCCGACCTGAAGAGCACCCAGAACGCCATCGACAAG
ATCACCAACAAGGTGAACAGCGTGATCGAGAAGATGAACACCCAGTTCACCGCCGTGGGCAAGGAGT
TCAACCACCTGGAGAAGCGCATCGAGAACCTGAACAAGAAGGTGGACGACGGCTTCCTGGACGTGT
GGACCTACAACGCCGAGCTGCTGGTGCTGCTGGAGAACGAGCGCACCCTGGACTACCACGACAGC
AACGTGAAGAACCTGTACGAGAAGGTGCGCCACCAGCTGAAGAACAACGCCAAGGAGATCGGCAAC
GGCTGCTTCGAGTTCTACCACAAGTGCGACAACACCTGCATGGAGAGCGTGAAGAACGGCACCTACG
ACTACCCCAAGTACAGCGAGGAGGCCAAGCTGAACCGCGAGAAGATCGACGGCGTGAAGCTGGAC
AGCACCCGCATCTACCAGATCCTGGCCATCTACAGCACCGTGGCCAGCAGCCTGGTGCTGGTGGTG
AGCCTGGGCGCCATCAGCTTCTGGATGTGCAGCAACGGCAGCCTGCAGTGCCGCATCTGCATCTtga

**Strain:** A/Wisconsin/67/2022
**Codon Optimization Scheme:** 2
**Source:** 2
ATGAAGGCCATTCTGGTGGTGATGCTGTACACCTTTACAACCGCCAATGCCGACACCCTGTGTATCGG
ATACCACGCCAACAACAGCACCGACACCGTGGACACCGTGCTGGAGAAGAACGTCACCGTGACCCA
CTCTGTGAACCTCCTGGAAGATAAGCACAACGGAAAGCTGTGCAAGCTGAGAGGCGTGGCCCCTCTG
CACCTGGGACAGTGCAACATCGCCGGCTGGATCCTGGGCAACCCTGAGTGCGAGAGCCTGAGCAC
AGCCAGAAGCTGGTCCTATATCGTGGAAACCAGCAACAGCGACAATGGCACCTGCTACCCTGGAGAT
TTCATCAATTACGAGGAACTGCGGGAACAGCTGAGCAGCGTGTCCAGCTTCGAGAGATTCGAGATCTT
CCCCAAGACCTCCAGCTGGCCCAACCACGACTCCGATAACGGCGTGACCGCCGCCTGCAGCCACG
CCGGAGCTAGATCTTTTTACAAGAATCTGATCTGGCTGGTAAAAAAAGGCAAGTCCTACCCAAAGATTAA
CCAGACCTACATCAACGACAAGGGCAAAGAGGTGCTGGTTCTGTGGGGCATCCATCACCCCCCCAC

AATCACCGACCAAGAGAGCCTGTACCAGAATGCCGACGCCTACGTGTTCGTGGGCACAAGTAGATAC
AGCAAGAAATTCAAGCCCGAGATCGCCACAAGACCTAAGGTCAGAGATCAGGCCGGCAGAATGAACT
ACTACTGGACCCTGGTTGAGCCTGGCGACAAGATCACCTTCGAGGCCACCGGCAATCTGGTGGCCC
CTCGGTACGCCTTCACCATGGAAAAGGAAGCTGGATCTGGCATCATCATTTCTGACACACCTGTGCAC
GATTGCAACGCCACATGCCAGACACCAGAGGGCGCTATCAACACCTCTCTGCCTTTCCAGAACGTGC
ACCCTATCACCATCGGCAAGTGCCCTAAGTACGTGCGGAGCACAAAGCTGAGGCTGGCCACTGGAC
TGCGGAACGTGCCCAGCATCCAAAGCCGCGGCCTGTTCGGAGCCATCGCCGGCTTCATCGAGGGC
GGATGGACAGGTATGGTGGATGGCTGGTACGGCTACCACCACCAGAACGACCAGGGCAGCGGCTAC
GCCGCTGATCTCAAGAGCACACAGAATGCTATTGACAAAATCACAAACAAAGTGAACTCCGTGATCGA
GAAAATGAACACACAGTTCACCGCCGTGGGCAAAGAGTTCAACCACCTGGAAAAAGAATCGAGAAC
CTGAACAAGAAGGTGGACGACGGCTTTCTGGACGTGTGGACCTATAACGCTGAGCTGCTGGTGTTGCT
GGAAAACGAGCGGACCCTGGATTACCACGACAGCAACGTGAAGAACCTGTATGAGAAGGTGCGGCA
CCAGCTGAAAAACAACGCAAAGGAAATCGGCAACGGTTGTTTTGAATTCTACCATAAGTGCGACAACAC
CTGTATGGAATCTGTGAAGAACGGCACATACGACTATCCTAAGTACTCCGAGGAAGCCAAGCTGAATA
GAGAGAAGATCGATGGAGTGAAGCTCGACTCTACCAGAATCTACCAAATCCTGGCTATCTACAGCACC
GTGGCCAGCAGCCTTGTCCTGGTGGTCAGCCTCGGCGCCATCAGCTTCTGGATGTGTAGCAATGGCA
GCCTGCAGTGTAGAATCTGCATCtga

**Strain:** A/Wisconsin/67/2022
**Codon Optimization Scheme:** 3
**Source:** 1*, 2, 3
*Underlined C is replaced with a G
ATGAAGGCCATCCTGGTGGTGATGCTGTACACCTTCACCACCGCCAACGCCGACACCCTGTGCATCG
GCTACCACGCCAACAACAGCACCGACACCGTGGACACCGTGCTGGAGAAGAACGTGACCGTGACC
CACAGCGTGAACCTGCTGGAGGACAAGCACAACGGCAAGCTGTGCAAGCTGAGAGGCGTGGCCCC
CCTGCACCTGGGGCAGTGCAACATCGCCGGCTGGATCCTGGGCAACCCCGAGTGCGAGAGCCTGA
GCACCGCTAGAAGCTGGAGCTACATCGTGGAGACCTCCAATAGTGACAATGGCACCTGTTACCCCGG
CGACTTCATCAACTACGAGGAGCTGAGAGAGCAGCTGAGCAGCGTGAGCAGCTTCGAGAGATTCGAG
ATCTTCCCCAAGACAAGCAGCTGGCCCAACCACGACAGTGACAATGGAGTGACGGCCGCCTGCAGC
CACGCCGGCGCTAGAAGCTTCTACAAGAACCTGATCTGGCTGGTGAAGAAGGGCAAGAGCTACCCC
AAGATCAATCAGACCTACATCAACGACAAGGGCAAAGAGGTGCTGGTGCTGTGGGGCATCCACCACC
CCCCCACCATCACCGACCAAGAGAGCCTGTATCAGAACGCCGACGCCTACGTGTTCGTGGGCACAA
GCAGATACAGCAAGAAGTTCAAGCCCGAGATCGCCACAAGACCCAAGGTGAGAGACCAAGCCGGCA
GAATGAACTACTACTGGACCCTGGTGGAGCCCGGCGACAAGATCACCTTCGAGGCCACCGGCAACC
TGGTGGCCCCTAGATACGCCTTCACCATGGAGAAGGAGGCCGGCAGCGGCATCATCATCAGCGACA
CCCCCGTGCACGACTGCAACGCCACCTGTCAGACCCCCGAGGGCGCCATCAACACAAGCCTGCCC
TTTCAGAACGTGCACCCCATCACCATCGGCAAGTGCCCCAAGTACGTGAGAAGCACCAAGCTGAGAC
TGGCCACCGGCCTGAGAAACGTGCCTAGCATTCAGAGCAGAGGCCTGTTCGGCGCCATCGCCGGCT
TCATCGAGGGCGGCTGGACCGGCATGGTGGACGGCTGGTACGGCTACCACCATCAGAACGACCAA
GGCAGCGGCTACGCCGCCGAC**CTC***AAGAGCACACAGAACGCCATCGATAAGATCACGAACAAGGT
GAACAGCGTGATCGAGAAGATGAACACACAGTTCACCGCCGTGGGCAAGGAGTTCAACCACCTGGA
GAAGAGAATCGAGAACCTGAACAAGAAGGTGGACGACGGCTTCCTGGACGTGTGGACCTACAACGC
CGAGCTGCTGGTGCTGCTGGAGAACGAGAGAACCCTGGACTACCACGACAGCAACGTGAAGAACCT
GTACGAGAAGGTGAGACATCAGCTGAAGAACAACGCCAAGGAGATCGGCAACGGCTGCTTCGAGTTC
TACCACAAGTGCGACAACACCTGCATGGAGAGCGTGAAGAACGGCACCTACGACTACCCCAAGTAC
AGCGAGGAGGCCAAGCTGAACAGAGAGAAGATCGACGGCGTGAAGCTGGACAGCACAAGAATCTAT

CAGATCCTGGCCATCTACAGCACCGTGGCTAGCAGCCTGGTGCTGGTGGTGAGCCTGGGCGCCATC
AGCTTCTGGATGTGCAGCAACGGCAGCCTGCAGTGCAGAATCTGCATCtga

**Strain:** A/Sydney/5/2021
**Codon Optimization Scheme:** 2
**Source:** 2
ATGAAGGCCATCCTGGTCGTGATGCTGTACACCTTTACCACAGCCAATGCCGACACACTGTGCATCGG
CTATCACGCCAACAATAGCACCGATACCGTGGACACTGTGCTGGAAAAGAATGTGACCGTGACCCAC
AGCGTGAATCTGCTGGAAGATAAGCACAACGGAAAGCTCTGCAAGCTGCGGGGCGTCGCCCCTCTG
CATCTGGGCCAGTGCAACATCGCCGGCTGGATCTTAGGCAACCCTGAGTGCGAGTCCCTGAGCACA
GCCCGGAGCTGGTCCTACATCGTGGAAACCTCTAACTCTGATAACGGCACATGTTACCCCGGCAATTT
CATCAACTACGAGGAACTGAGAGAGCAGCTGAGCTCCGTGTCTTCTTTCGAGCGATTTGAGATCTTCC
CTAAAACCAGCTCTTGGCCTAACCACGACAGCGACAACGGCGTGACAGCTGCATGCCCCCACGCCG
GCGCCAAAAGCTTTTACAAGAACCTGATCTGGCTGGTCAAAAAGGGCAAAAGTTATCCAAAGATTAATC
AGACCTACATCAACGACAAGGGCAAGGAAGTGCTGGTGCTGTGGGGCATCCACCACCCCCCCACAA
TCACAGATCAGGAGAGCCTGTATCAGAACGCTGATGCCTACGTGTTCGTGGGCACCAGCCGGTACAG
CAAGAAGTTCAAGCCCGAGATCGCCGCTAGACCTAAGGTGCGGGACAGAGCCGGAAGAATGAACTA
CTACTGGACCCTGGTGGAACCAGGAGATAAGATCACCTTCGAGGCCACAGGCAACCTTGTTGCTCCT
AGATACGCCTTCACCATGGAAAAGGACGCCGGCAGCGGCATCATCATCTCAGATACCCCAGTGCAC
GATTGCAATACCACCTGTCAGACCCCTGAGGGCGCTATCAATACCAGCCTGCCTTTCCAGAACGTGC
ACCCTATCACCATTGGAAAATGCCCTAAGTACGTGCGCAGCACCAAGCTGCGGCTGGCCACCGGACT
GAGAAACGTGCCTAGCATCCAGAGCAGAGGCCTGTTCGGCGCCATTGCCGGATTTATCGAGGGCGG
CTGGACAGGCATGGTGGACGGCTGGTACGGTTACCACCACCAGAATGAGCAAGGCTCTGGCTACGC
CGCCGACCTGAAGTCTACACAAAACGCCATTGACAAGATCACGAACAAGGTGAATTCTGTGATCGAGA
AGATGAACACCCAGTTCACAGCTGTTGGAAAGGAGTTCAACCACCTTGAGAAAAGAATCGAGAACTTG
AACAAGAAAGTCGACGACGGCTTCCTGGATATCTGGACCTACAACGCCGAGCTGCTGGTGCTGCTGG
AAAACGAGAGAACACTGGACTACCACGACAGCAATGTGAAGAACCTGTACGAGAAGGTGCGGAACCA
ACTGAAGAACAACGCCAAGGAAATCGGCAACGGCTGCTTCGAGTTCTACCATAAGTGCGACAACACC
TGTATGGAAAGCGTGAAGAACGGCACCTACGACTACCCCAAATACAGTGAAGAGGCCAAGCTGAACA
GAGAAAAAATCGACGGCGTGAAGCTGGACAGCACAAGAATCTACCAGATCCTGGCTATCTACAGCAC
CGTGGCCAGCAGCCTGGTCCTGGTGGTGTCCCTGGGCGCCATCAGCTTCTGGATGTGCAGCAACGG
AAGCCTGCAGTGTAGAATCTGCATCtga

**Strain:** A/Wisconsin/588/2019
**Codon Optimization Scheme:** 2
**Source:** 2
ATGAAGGCCATTCTGGTTGTGATGCTGTACACCTTTACAACAGCCAATGCAGACACCCTGTGCATCGG
CTACCACGCCAACAACAGCACCGACACCGTGGACACAGTGCTGGAAAAGAACGTGACCGTGACCCA
CAGCGTTAATCTGCTGGAAGATAAGCACAACGGCAAACTGTGTAAACTGAGAGGAGTGGCTCCTCTGC
ACCTGGGCAAGTGCAACATTGCCGGATGGATCCTGGGAAACCCCGAGTGTGAATCTCTCAGCACCGC
CAGAAGCTGGTCTTATATCGTGGAAACCTCTAACAGCGACAACGGCACCTGCTACCCCGGCGACTTC
ATCAACTACGAGGAACTGAGAGAGCAGCTGAGCAGCGTGTCCAGTTTCGAGCGGTTCGAGATCTTCC
CAAAAACCAGCTCCTGGCCTAACCACGATAGCGATAATGGCGTGACAGCTGCCTGCCCCCACGCCG
GTGCCAAGAGCTTCTACAAGAACCTGATCTGGCTGGTGAAAAAGGGCAAGTCCTACCCTAAGATCAAT
CAAACCTATATTAACGACAAAGGCAAGGAGGTGCTCGTGCTTTGGGGCATCCACCACCCCCCCACCA

TCGCCGATCAGCAGAGCCTGTACCAGAACGCCGACGCCTACGTGTTCGTGGGAACATCCCGGTACA
GCAAGAAATTCAAGCCTGAGATCGCCACAAGACCTAAGGTGCGGGACCAGGAGGGCAGAATGAACTA
CTACTGGACCCTGGTCGAGCCTGGCGATAAGATCACTTTTGAGGCTACCGGCAACCTCGTCGCCCCT
AGATACGCCTTCACCATGGAAAGAGATGCCGGCTCTGGCATCATCATCTCTGATACCCCTGTGCACGA
CTGCAATACCACCTGCCAGACCCCAGAGGGCGCCATCAATACGAGCCTGCCCTTCCAGAACGTTCAT
CCTATCACAATCGGCAAGTGTCCTAAGTACGTCAAGAGCACAAAGCTGCGGCTGGCTACAGGACTGA
GGAACGTGCCTTCTATCCAGAGCAGAGGCCTGTTCGGCGCTATCGCCGGCTTCATCGAGGGCGGCT
GGACAGGCATGGTGGACGGCTGGTACGGCTACCACCACCAGAACGAGCAGGGCTCTGGATACGCC
GCTGACCTGAAGTCTACACAGAATGCCATCGATAAGATCACCAACAAGGTGAACAGCGTGATCGAGAA
AATGAACACCCAGTTCACAGCCGTGGGCAAGGAATTCAACCACCTGGAAAAGAGAATCGAGAACCTG
AACAAGAAGGTGGACGACGGCTTTCTGGACATCTGGACCTACAACGCCGAGCTGCTGGTCCTGCTGG
AAAACGAGCGCACCCTGGACTACCATGACAGCAACGTGAAGAACCTGTACGAGAAAGTGAGAAACCA
GCTGAAGAACAATGCCAAGGAAATCGGGAACGGCTGCTTCGAGTTCTACCACAAGTGCGACAACACC
TGTATGGAATCCGTGAAGAATGGCACCTACGACTATCCTAAGTACAGCGAGGAAGCCAAGCTGAACCG
GGAAAAAATCGACGGCGTGAAGTTAGATAGCACCAGAATCTATCAAATCCTGGCTATCTACAGCACAGT
GGCCAGCAGTCTGGTGCTGGTGGTCAGCCTGGGAGCCATTAGCTTTTGGATGTGCAGCAACGGTAGC
CTGCAGTGCAGAATCTGCATCtga

**<u>Type/Subtype:</u>** <u>H3</u>

**Strain:** A/Darwin/6/2021
**Codon Optimization Scheme:** 1
**Source:** 3
ATGAAGACCATCATCGCCCTGAGCAACATCCTGTGCCTGGTGTTCGCCCAGAAGATCCCCGGCAACG
ACAACAGCACCGCCACCCTGTGCCTGGGCCACCACGCCGTGCCCAACGGCACCATCGTGAAGACC
ATCACCAACGACCGCATCGAGGTGACCAACGCCACCGAGCTGGTGCAGAACAGCAGCATCGGCGA
GATCTGCGGCAGCCCCCACCAGATCCTGGACGGCGGCAACTGCACCCTGATCGACGCCCTGCTGG
GCGACCCCCAGTGCGACGGCTTCCAGAACAAGGAGTGGGACCTGTTCGTGGAGCGCAGCCGCGCC
AACAGCAACTGCTACCCCTACGACGTGCCCGACTACGCCAGCCTGCGCAGCCTGGTGGCCAGCAG
CGGCACCCTGGAGTTCAAGAACGAGAGCTTCAACTGGACCGGCGTGAAGCAGAACGGCACCAGCA
GCGCCTGCATCCGCGGCAGCAGCAGCAGCTTCTTCAGCCGCCTGAACTGGCTGACCAGCCTGAACA
ACATCTACCCCGCCCAGAACGTGACCATGCCCAACAAGGAGCAGTTCGACAAGCTGTACATCTGGG
GCGTGCACCACCCCGACACCGACAAGAACCAGATCAGCCTGTTCGCCCAGAGCAGCGGCCGCATC
ACCGTGAGCACCAAGCGCAGCCAGCAGGCCGTGATCCCCAACATCGGCAGCCGCCCCCGCATCC
GCGACATCCCCAGCCGCATCAGCATCTACTGGACCATCGTGAAGCCCGGCGACATCCTGCTGATCA
ACAGCACCGGCAACCTGATCGCCCCCCGCGGCTACTTCAAGATCCGCAGCGGCAAGAGCAGCATC
ATGCGCAGCGACGCCCCCATCGGCAAGTGCAAGAGCGAGTGCATCACCCCCAACGGCAGCATCCC
CAACGACAAGCCCTTCCAGAACGTGAACCGCATCACCTACGGCGCCTGCCCCCGCTACGTGAAGCA
GAGCACCCTGAAGCTGGCCACCGGCATGCGCAACGTGCCCGAGAAGCAGACCCGCGGCATCTTCG
GCGCCATCGCCGGCTTCATCGAGAACGGCTGGGAGGGCATGGTGGACGGCTGGTACGGCTTCCGC
CACCAGAACAGCGAGGGCCGCGGCCAGGCCGCCGACCTGAAGAGCACCCAGGCCGCCATCGAC
CAGATCAACGGCAAGCTGAACCGCCTGATCGGCAAGACCAACGAGAAGTTCCACCAGATCGAGAAG
GAGTTCAGCGAGGTGGAGGGCCGCGTGCAGGACCTGGAGAAGTACGTGGAGGACACCAAGATCGA
CCTGTGGAGCTACAACGCCGAGCTGCTGGTGGCCCTGGAGAACCAGCACACCATCGACCTGACCGA
CAGCGAGATGAACAAGCTGTTCGAGAAGACCAAGAAGCAGCTGCGCGAGAACGCCGAGGACATGGG
CAACGGCTGCTTCAAGATCTACCACAAGTGCGACAACGCCTGCATCGGCAGCATCCGCAACGAGAC
CTACGACCACAACGTGTACCGCGACGAGGCCCTGAACAACCGCTTCCAGATCAAGGGCGTGGAGCT

GAAGAGCGGCTACAAGGACTGGATCCTGTGGATCAGCTTCGCCATGAGCTGCTTCCTGCTGTGCATC
GCCCTGCTGGGCTTCATCATGTGGGCCTGCCAGAAGGGCAACATCCGCTGCAACATCTGCATCtga

**Strain:** A/Darwin/6/2021
**Codon Optimization Scheme:** 2
**Source:** 1, 2, 3
ATGAAGACCATCATCGCCCTGAGCAACATCCTCTGCCTGGTGTTCGCTCAAAAGATCCCCGGCAACG
ACAATAGCACAGCCACCCTGTGTCTGGGCCACCATGCCGTGCCTAATGGCACCATCGTTAAGACCAT
CACCAACGATAGAATTGAAGTGACAAACGCTACAGAGCTGGTGCAAAATAGCTCCATCGGAGAGATCT
GCGGCTCCCCTCACCAGATCCTGGACGGCGGCAATTGCACCCTGATCGATGCCCTGCTGGGCGAC
CCTCAGTGTGATGGATTTCAGAACAAGGAATGGGACCTGTTTGTGGAACGCAGCAGAGCCAACAGCAA
CTGCTACCCCTACGACGTGCCCGACTACGCCTCTCTGAGATCTCTGGTGGCCAGCTCAGGCACCCT
GGAGTTCAAGAACGAGAGCTTCAACTGGACAGGCGTCAAGCAGAATGGCACCAGCAGCGCCTGCAT
ACGCGGAAGCAGCAGCAGTTTCTTCAGCAGACTGAACTGGCTGACAAGCCTGAACAACATTTACCCTG
CCCAGAACGTCACAATGCCTAACAAGGAACAGTTTGACAAGCTGTACATCTGGGGCGTGCACCACCC
TGATACCGACAAGAACCAGATCAGCCTGTTCGCCCAGTCTTCTGGCAGAATCACCGTGTCTACAAAAA
GATCCCAGCAGGCTGTGATCCCCAATATCGGTAGCAGACCCCGGATTCGGGACATCCCTAGCAGAAT
CAGCATCTATTGGACCATCGTGAAACCCGGAGATATCCTGCTGATCAACTCCACAGGCAACCTGATTG
CCCCAAGGGGCTACTTCAAGATCCGGAGCGGCAAGAGCAGCATCATGCGGAGCGACGCCCCTATC
GGAAAATGCAAGAGCGAGTGCATCACGCCTAACGGCTCCATCCCGAACGATAAGCCTTTCCAGAACG
TGAACAGAATCACCTACGGCGCTTGTCCTAGATATGTGAAGCAAAGCACTCTGAAACTGGCCACCGGC
ATGAGAAACGTTCCAGAGAAACAGACCAGAGGCATCTTCGGCGCTATCGCCGGGTTCATCGAGAACG
GCTGGGAAGGCATGGTGGACGGCTGGTACGGCTTCAGACACCAGAACTCTGAGGGCCGGGGCCAG
GCCGCTGATCTCAAGAGCACACAGGCCGCCATCGACCAGATCAACGGAAAGCTGAACCGGCTGATC
GGCAAAACCAATGAGAAGTTCCACCAGATCGAGAAGGAATTCTCCGAGGTGGAAGGCAGAGTGCAGG
ACCTGGAAAAGTACGTGGAGGACACCAAGATCGACCTGTGGAGCTACAACGCCGAGCTGCTGGTGG
CACTGGAGAACCAGCACACCATCGACCTCACCGATAGCGAGATGAACAAGCTGTTCGAGAAGACCAA
AAAGCAGCTGCGGGAAAATGCTGAAGATATGGGCAACGGATGTTTTAAGATCTACCACAAGTGCGACA
ACGCCTGCATCGGTTCTATCAGAAATGAGACATACGACCACAACGTGTACAGAGATGAGGCCCTGAAC
AACAGATTCCAAATCAAGGGCGTGGAACTGAAGTCTGGCTACAAGGACTGGATCCTGTGGATCAGTTTT
GCCATGAGCTGCTTCCTGCTGTGCATTGCCCTGCTCGGCTTCATCATGTGGGCCTGTCAGAAAGGCAA
CATCCGGTGCAACATCTGCATCtga

**Strain:** A/Darwin/6/2021
**Codon Optimization Scheme:** 3
**Source:** 1, 3
ATGAAAACCATCATCGCCCTGAGCAACATCCTGTGCCTGGTTTTCGCGCAGAAAATTCCCGGCAACGA
CAACAGCACCGCCACCCTGTGCCTGGGCCACCACGCCGTGCCCAACGGCACCATCGTGAAAACCA
TAACCAACGACAGAATCGAGGTGACCAACGCCACCGAGCTGGTGCAGAACAGCAGCATCGGCGAGA
TCTGCGGCAGCCCCCATCAGATCCTGGACGGCGGCAACTGCACCCTGATCGACGCGCTGCTGGGC
GACCCTCAGTGCGACGGCTTTCAGAACAAGGAGTGGGACCTGTTCGTGGAGAGAAGCAGAGCCAAC
AGCAACTGCTACCCCTATGACGTGCCGGACTACGCTAGCCTGAGAAGCCTGGTGGCTAGCAGCGGC
ACCCTGGAGTTCAAGAACGAGAGCTTCAACTGGACCGGCGTGAAGCAGAACGGCACAAGCAGCGCC
TGCATCAGAGGCAGCAGCAGCAGCTTCTTCAGCAGACTGAACTGGCTGACAAGCCTGAACAACATCT
ACCCCGCTCAGAACGTGACCATGCCCAACAAGGAGCAGTTCGACAAGCTGTACATCTGGGGCGTGC
ACCACCCCGACACCGACAAGAATCAGATCAGCCTGTTCGCTCAGAGTAGTGGCAGAATCACCGTGAG
CACCAAGAGATCTCAGCAAGCCGTGATCCCCAACATCGGCAGCAGACCTAGAATCAGAGACATCCCT

AGCAGAATCAGCATCTACTGGACTATCGTGAAGCCCGGCGACATCCTGCTGATCAACAGCACCGGCA
ACCTGATCGCCCCTAGAGGCTACTTCAAGATCAGAAGCGGCAAGAGCAGCATCATGAGAAGCGACG
CCCCCATCGGCAAGTGCAAGAGCGAGTGCATCACCCCCAACGGTAGCATACCCAATGATAAGCCCTT
TCAGAACGTGAACAGAATCACCTACGGCGCCTGCCCTAGATACGTGAAGCAGAGCACCCTGAAGCTG
GCCACCGGCATGAGAAACGTGCCCGAGAAGCAGACAAGAGGCATCTTCGGCGCCATCGCCGGCTT
CATCGAGAACGGCTGGGAGGGCATGGTGGACGGCTGGTACGGCTTCAGACATCAGAACAGCGAGG
GCAGAGGCCAAGCCGCCGACCTGAAGAGCACCCAAGCCGCCATCGATCAGATCAACGGCAAGCTG
AACAGACTGATCGGCAAGACCAACGAGAAGTTCCATCAGATCGAGAAGGAGTTCAGCGAGGTGGAGG
GCAGAGTGCAAGACCTGGAGAAGTACGTGGAGGACACCAAGATCGACCTGTGGAGCTACAACGCCG
AGCTGCTGGTGGCCCTGGAGAATCAGCACACCATCGACCTGACCGACAGCGAGATGAACAAGCTGTT
CGAGAAGACCAAGAAGCAGCTGAGAGAGAACGCCGAGGACATGGGCAACGGCTGCTTCAAGATCTA
CCACAAGTGCGACAACGCCTGCATCGGCAGCATCAGAAACGAGACCTACGACCACAACGTGTACAG
AGACGAGGCCCTGAACAACAGATTTCAGATCAAGGGCGTGGAGCTGAAGAGCGGCTACAAGGACTG
GATCCTGTGGATCAGCTTCGCCATGAGCTGCTTCCTGCTGTGCATCGCTCTGCTGGGCTTCATCATGT
GGGCCTGTCAGAAGGGCAACATCAGATGCAACATCTGCATCtga

**Strain:** A/Cambodia/e826360/2020
**Codon Optimization Scheme:** 2
**Source:** 2
ATGAAGACCATCATCGCCTTGTCTTACATCCTGTGCCTGGTGTTCGCCCAGAAAATCCCCGGCAACGA
CAACTCTACCGCCACACTGTGTCTGGGCCACCACGCCGTGCCCAATGGAACAATCGTGAAAACAATC
ACAAACGATAGAATCGAGGTGACCAACGCCACCGAGCTCGTGCAAAATAGCAGCATCGGCGAGATCT
GCGATAGCCCTCACCAGATCCTGGACGGTGGCAACTGCACCCTGATCGACGCCCTGCTCGGCGATC
CTCAGTGCGACGGCTTTCAGAACAAGGAATGGGACCTGTTCGTGGAAAGAAGCAGAGCCAACAGCAA
CTGCTACCCCTACGACGTGCCAGACTACGCCTCCCTGCGGAGCCTGGTCGCCAGCAGCGGCACCC
TGGAATTCAAGAACGAAAGCTTCAACTGGACCGGAGTTAAGCAGAACGGCACCAGCTCAGCCTGTATA
AGAGGCTCCTCTAGCAGCTTCTTCTCCAGACTGAATTGGCTTACACACCTGAACTACAAGTACCCTGC
CCTGAATGTGACCATGCCTAACAACGAGCAGTTCGACAAGCTGTACATCTGGGGCGTGCACCATCCT
CGGACCGACAAAGACCAGATCAGCCTGTTTGCCCAGCCTTCTGGCAGAATCACCGTGAGCACCAAAA
GAAGCCAGCAGGCCGTCATCCCCAATATCGGATCTAGACCTAGGATCCGCGACATCCCAAGCAGGA
TCAGCATTTACTGGACAATCGTGAAGCCTGGCGACATCCTGCTGATTAACAGCACCGGCAACCTGATC
GCCCCTCGGGGCTACTTCAAGATCCGCTCCGGTAAGAGCTCCATCATGCGGAGTGATGCTCCTATCG
GCAAATGCAAGTCCGAGTGCATCACACCTAACGGCTCTATCCCTAACGACAAGCCCTTCCAAAATGTG
AACCGGATCACCTATGGCGCTTGCCCCAGATACGTGAAGCAGAGCACCCTGAAACTGGCCACAGGA
ATGAGAAACGTTCCCGAGAAGCAAACAAGAGGCATCTTCGGCGCCATCGCCGGCTTCATCGAGAACG
GCTGGGAGGGCATGGTGGACGGCTGGTACGGCTTTAGACACCAGAACAGCGAGGGCAGAGGCCAG
GCCGCTGACCTCAAGAGCACACAGGCTGCTATCGATCAGATCAACGGCAAGCTGAATCGGCTGATCG
GCAAGACCAACGAGAAGTTCCACCAGATAGAGAAAGAGTTCAGCGAGGTGGAAGGAAGAGTGCAGG
ACCTGGAAAAATATGTGGAGGACACCAAGATCGATCTGTGGAGCTACAACGCCGAGCTGCTGGTGGC
CCTGGAAAACCAGCACACCATCGACCTGACCGACAGCGAAATGAACAAGCTGTTCGAGAAGACCAA
GAAGCAGCTGCGGGAAAATGCTGAAGATATGGGAAATGGATGTTTTAAGATCTACCACAAGTGCGACAA
CGCTTGCATCGGCAGCATCAGAAACGAGACATACGATCACAACGTGTACAGAGATGAGGCCCTGAAC
AACAGATTCCAGATTAAGGGCGTGGAACTGAAGTCTGGATACAAGGACTGGATCCTGTGGATCAGCTTT
GCCATGTCTTGTTTCCTGCTGTGTATCGCCCTGCTGGGCTTCATTATGTGGGCCTGCCAAAAGGGCAA
CATCCGGTGCAACATCTGCATCtga

**Strain:** A/HongKong/45/2019

**Codon Optimization Scheme:** 2
**Source:** 2
ATGAAGACCATCATTGCCCTGAGCTACATCCTGTGCCTGGGCTTCACCCAAAAGATCCCCGGCAATGA
CAACAGCACAGCCACACTGTGCCTGGGCCACCACGCCGTGCCAAATGGAACAATCGTGAAGACCAT
CACCAATGATAGAATCGAGGTGACCAACGCTACAGAGCTGGTCCAGAACAGCAGTATCGGCGAGATC
TGTGATTCTCCTCACCAGATCCTGGACGGCGGCAACTGCACCCTGATTGATGCCCTGCTGGGCGACC
CCCAGTGCGACGGCTTCCAGAACAAAAAGTGGGACCTCTTCGTGGAAAGAAGTCGGGCCTACTCTAA
TTGCTACCCTTACGACGTGCCTGACTACGCCAGCCTGAGAAGCCTGGTGGCCAGCTCTGGAACCCTG
GAATTCAAGAACGAATCTTTTAACTGGGCCGGCGTGACACAAAACGGCAAGAGCTTCTCTTGCATCAG
AGGCAGCAGCAGCAGCTTCTTCAGCAGACTGAACTGGCTGACCCACCTGAATTACACCTACCCCGCC
CTGAACGTGACCATGCCTAACAAGGAACAGTTCGACAAGCTGTATATCTGGGGCGTGCACCATCCTGG
CACAGACAAAGACCAGATCTCTCTGTACGCTCAGAGCAGCGGCAGAATCACAGTGTCCACCAAACGG
TCCCAGCAGGCTGTGATCCCTAACATCGGTTCTCGTCCAAGAATCAGAGATATCCCCAGCAGAATCAG
CATTTATTGGACCATCGTGAAGCCCGGAGATATCCTGCTTATTAACAGCACCGGCAATCTGATCGCCC
CTAGGGGCTACTTCAAGATCCGGAGCGGCAAGTCCAGCATCATGAGAAGCGACGCCCCTATCGGGA
AATGTAAAAGCGAGTGCATCACCCCTAATGGCAGCATCCCAAACGACAAACCTTTCCAAAACGTGAAC
AGAATAACATACGGCGCTTGTCCTAGATACGTGAAACAGAACACCCTGAAGCTGGCCACAGGCATGC
GGAACGTGCCCGAGAAACAGACCAGAGGAATTTTCGGAGCCATCGCCGGCTTTATCGAGAACGGCTG
GGAGGGCATGGTGGACGGCTGGTACGGCTTTAGACACCAGAACTCTGAGGGCCGCGGACAGGCTG
CCGATCTGAAAAGCACCCAAGCCGCTATCGATCAGATCAACGGCAAGCTGAACCGGCTGATCGGCA
AGACCAACGAGAAGTTCCACCAGATCGAGAAGGAATTTTCTGAAGTGGAAGGCAGAGTGCAGGACCT
GGAAAAGTACGTTGAGGACACCAAGATCGACCTGTGGAGCTACAACGCCGAACTGCTGGTGGCTCTG
GAAAACCAGCACACCATCGACCTGACCGACAGCGAGATGAACAAGCTGTTCGAGAAGACAAAGAAG
CAGCTGCGGGAAAACGCCGAGGATATGGGCAACGGATGTTTTAAGATCTACCACAAGTGCGATAATGC
CTGTATCGGCTCCATCCGAAATGAGACATATGACCACAACGTGTACCGGGACGAGGCCCTGAACAAC
CGGTTCCAGATCAAGGGCGTCGAGCTGAAGTCCGGCTACAAGGATTGGATCCTGTGGATCAGCTTCG
CCATCAGCTGCTTCCTGCTGTGCGTGGCCCTCCTCGGATTCATCATGTGGGCCTGCCAGAAGGGCAA
CATCAGATGCAACATCTGCATCtga

**Type/Subtype:** B/Y

**Strain:** B/Phuket/3073/2013
**Codon Optimization Scheme:** 1
**Source:** 3
ATGAAGGCCATCATCGTGCTGCTGATGGTGGTGACCAGCAACGCCGACCGCATCTGCACCGGCATC
ACCAGCAGCAACAGCCCCCACGTGGTGAAGACCGCCACCCAGGGCGAGGTGAACGTGACCGGCG
TGATCCCCCTGACCACCACCCCCACCAAGAGCTACTTCGCCAACCTGAAGGGCACCCGCACCCGC
GGCAAGCTGTGCCCCGACTGCCTGAACTGCACCGACCTGGACGTGGCCCTGGGCCGCCCCATGTG
CGTGGGCACCACCCCCAGCGCCAAGGCCAGCATCCTGCACGAGGTGCGCCCCGTGACCAGCGGC
TGCTTCCCCATCATGCACGACCGCACCAAGATCCGCCAGCTGCCCAACCTGCTGCGCGGCTACGAG
AAGATCCGCCTGAGCACCCAGAACGTGATCGACGCCGAGAAGGCCCCCGGCGGCCCCTACCGCC
TGGGCACCAGCGGCAGCTGCCCCAACGCCACCAGCAAGATCGGCTTCTTCGCCACCATGGCCTGG
GCCGTGCCCAAGGACAACTACAAGAACGCCACCAACCCCCTGACCGTGGAGGTGCCCTACATCTGC
ACCGAGGGCGAGGACCAGATCACCGTGTGGGGCTTCCACAGCGACAACAAGACCCAGATGAAGAG
CCTGTACGGCGACAGCAACCCCCAGAAGTTCACCAGCAGCGCCAACGGCGTGACCACCCACTACG
TGAGCCAGATCGGCGACTTCCCCGACCAGACCGAGGACGGCGGCCTGCCCCAGAGCGGCCGCAT
CGTGGTGGACTACATGATGCAGAAGCCCGGCAAGACCGGCACCATCGTGTACCAGCGCGGCGTGCT

GCTGCCCCAGAAGGTGTGGTGCGCCAGCGGCCGCAGCAAGGTGATCAAGGGCAGCCTGCCCCTGA
TCGGCGAGGCCGACTGCCTGCACGAGGAGTACGGCGGCCTGAACAAGAGCAAGCCCTACTACACC
GGCAAGCACGCCAAGGCCATCGGCAACTGCCCCATCTGGGTGAAGACCCCCCTGAAGCTGGCCAA
CGGCACCAAGTACCGCCCCCCCGCCAAGCTGCTGAAGGAGCGCGGCTTCTTCGGCGCCATCGCC
GGCTTCCTGGAGGGCGGCTGGGAGGGCATGATCGCCGGCTGGCACGGCTACACCAGCCACGGCG
CCCACGGCGTGGCCGTGGCCGCCGACCTGAAGAGCACCCAGGAGGCCATCAACAAGATCACCAAG
AACCTGAACAGCCTGAGCGAGCTGGAGGTGAAGAACCTGCAGCGCCTGAGCGGCGCCATGGACGA
GCTGCACAACGAGATCCTGGAGCTGGACGAGAAGGTGGACGACCTGCGCGCCGACACCATCAGCA
GCCAGATCGAGCTGGCCGTGCTGCTGAGCAACGAGGGCATCATCAACAGCGAGGACGAGCACCTG
CTGGCCCTGGAGCGCAAGCTGAAGAAGATGCTGGGCCCCAGCGCCGTGGACATCGGCAACGGCTG
CTTCGAGACCAAGCACAAGTGCAACCAGACCTGCCTGGACCGCATCGCCGCCGGCACCTTCAACG
CCGGCGAGTTCAGCCTGCCCACCTTCGACAGCCTGAACATCACCGCCGCCAGCCTGAACGACGAC
GGCCTGGACAACCACACCATCCTGCTGTACTACAGCACCGCCGCCAGCAGCCTGGCCGTGACCCT
GATGCTGGCCATCTTCATCGTGTACATGGTGAGCCGCGACAACGTGAGCTGCAGCATCTGCCTGtga

**Strain:** B/Phuket/3073/2013
**Codon Optimization Scheme:** 2
**Source:** 3
ATGAAGGCCATCATCGTGCTGCTGATGGTCGTGACCTCTAACGCCGATAGAATCTGTACCGGCATCAC
CAGCAGCAACAGCCCCCACGTGGTGAAGACCGCCACCCAGGGCGAGGTGAACGTCACCGGCGTTA
TTCCTCTGACCACCACACCCACAAAGAGTTACTTCGCCAACCTGAAGGGAACCCGGACCAGAGGAAA
GCTGTGCCCTGATTGCCTTAACTGTACAGATCTGGACGTGGCCCTCGGCAGACCTATGTGCGTGGGC
ACCACCCCTAGCGCCAAGGCTAGCATCCTGCATGAGGTGAGACCCGTGACAAGCGGATGTTTTCCTA
TCATGCACGACAGAACCAAGATCAGACAGCTGCCTAACCTGCTGAGAGGCTACGAGAAGATCCGGCT
GAGCACTCAGAACGTGATCGACGCCGAGAAGGCCCCTGGCGGACCTTACCGGCTGGGCACATCCG
GCTCATGCCCCAATGCCACCTCTAAGATCGGCTTCTTCGCCACAATGGCCTGGGCTGTGCCCAAGGA
CAATTACAAGAACGCCACCAACCCCCTAACAGTGGAAGTGCCTTATATATGTACCGAGGGCGAGGAC
CAGATCACCGTGTGGGGCTTCCACAGCGACAACAAGACCCAAATGAAATCCCTGTATGGCGATAGCA
ACCCACAGAAATTCACCAGCTCTGCCAACGGCGTGACCACCCACTACGTGTCCCAGATCGGCGACTT
CCCAGATCAGACCGAAGATGGCGGCCTGCCCCAGAGCGGCCGGATCGTGGTGGACTACATGATGC
AGAAGCCTGGAAAGACAGGTACCATCGTGTACCAGCGGGGCGTGCTGCTGCCCCAAAAAGTCTGGT
GCGCCAGCGGCAGAAGCAAGGTGATTAAGGGCAGCCTGCCTCTGATCGGGGAAGCCGACTGTCTGC
ACGAGGAATACGGCGGCCTGAATAAGTCCAAGCCTTACTACACAGGCAAACATGCCAAGGCCATCGG
AAACTGCCCTATCTGGGTGAAAACCCCTCTGAAACTGGCCAACGGCACCAAGTACAGACCACCTGCC
AAGCTGCTGAAGGAACGGGGCTTTTTCGGCGCCATCGCTGGCTTTCTGGAAGGAGGTTGGGAGGGCA
TGATTGCCGGATGGCACGGATACACCTCCCACGGCGCTCACGGCGTGGCCGTTGCCGCTGATCTCA
AAAGCACTCAAGAGGCCATCAACAAAATCACAAAGAATCTGAATTCTCTGAGCGAACTGGAAGTGAAG
AACCTGCAGAGACTGTCTGGCGCAATGGACGAGCTGCACAACGAGATCCTGGAACTGGACGAGAAA
GTGGACGATCTGAGGGCTGATACCATCAGCAGCCAGATCGAGCTGGCCGTGCTGCTGTCCAATGAGG
GAATCATCAACAGCGAAGATGAGCACCTGCTGGCTCTGGAAAGAAAGCTGAAAAAGATGCTGGGCCC
TAGCGCCGTGGACATCGGCAACGGCTGCTTCGAGACAAAGCACAAGTGCAATCAGACATGCCTCGA
CAGAATCGCCGCCGGCACCTTTAACGCCGGCGAGTTCAGCCTGCCTACATTCGACAGCCTGAACATC
ACAGCCGCCAGCCTGAACGACGACGGCCTGGACAACCACACAATCCTGCTGTACTACAGCACCGC
CGCTTCTTCTCTGGCTGTGACACTGATGCTGGCCATCTTCATCGTGTACATGGTGTCGCGCGACAACGT
GTCTTGCAGCATCTGCCTGtga

**Strain:** B/Phuket/3073/2013
**Codon Optimization Scheme:** 3
**Source:** 1, 2, 3
ATGAAGGCCATCATCGTGCTGCTGATGGTGGTGACAAGCAACGCCGACAGAATCTGCACCGGCATCA
CAAGCAGCAACAGCCCCCACGTGGTGAAGACCGCCACCCAAGGCGAGGTGAACGTGACCGGCGTG
ATCCCCCTGACCACCACCCCCACCAAGAGCTACTTCGCCAACCTGAAGGGCACAAGAACAAGAGGC
AAGCTGTGCCCCGACTGCCTGAACTGCACGGACCTGGACGTGGCCCTGGGCAGACCCATGTGTGTG
GGTACCACGCCTAGCGCCAAGGCTAGCATCCTGCACGAGGTGAGACCCGTGACAAGCGGCTGCTTC
CCCATCATGCACGACAGAACCAAGATCAGACAGCTGCCCAACCTGCTGAGAGGCTACGAGAAGATC
AGACTGTCTACACAGAACGTGATAGATGCCGAGAAGGCACCCGGTGGACCCTACAGACTGGGCACA
AGCGGCAGCTGCCCCAACGCCACAAGCAAGATCGGCTTCTTCGCCACCATGGCCTGGGCCGTGCC
CAAGGACAACTACAAGAACGCCACCAACCCCCTGACCGTGGAGGTGCCCTACATCTGCACCGAGGG
CGAGGATCAGATCACCGTGTGGGGCTTCCACAGCGACAACAAGACACAGATGAAGAGTCTGTACGGC
GACAGCAACCCTCAGAAGTTCACAAGCAGCGCCAACGGCGTGACCACCCACTACGTGTCTCAGATC
GGCGACTTCCCCGATCAGACCGAGGACGGCGGCCTGCCTCAGAGCGGCAGAATCGTGGTGGACTA
CATGATGCAGAAGCCCGGCAAGACCGGCACCATCGTGTATCAGAGAGGCGTGCTGCTGCCTCAGAA
GGTGTGGTGCGCTAGCGGCAGAAGCAAGGTGATCAAGGGCAGCCTGCCCCTGATCGGCGAGGCCG
ACTGCCTGCACGAGGAGTACGGCGGCCTGAACAAGAGCAAGCCCTACTACACCGGCAAGCACGCC
AAGGCCATCGGCAACTGCCCCATCTGGGTGAAGACCCCCCTGAAGCTGGCCAACGGCACCAAGTAC
AGACCCCCCGCCAAGCTGCTGAAGGAGAGAGGCTTCTTCGGAGCCATCGCGGGTTTCCTCGAGGGG
GGGTGGGAAGGAATGATCGCCGGGTGGCATGGGTATACAAGCCACGGTGCGCATGGAGTGGCCGTG
GCCGCCGACCTCAAGAGCACCCAAGAGGCCATCAACAAGATCACCAAGAACCTGAACAGCCTGAGC
GAGCTGGAGGTGAAGAACCTGCAGAGACTGAGCGGCGCCATGGACGAGCTGCACAACGAGATCCTG
GAGCTGGACGAGAAGGTGGACGACCTGAGAGCCGACACCATCAGCTCTCAGATCGAGCTGGCCGTG
CTGCTGAGCAACGAGGGCATCATCAACAGCGAGGACGAGCACCTGCTGGCCCTGGAGAGAAAGCTG
AAGAAGATGCTGGGCCCTAGCGCCGTGGACATCGGCAACGGCTGCTTCGAGACCAAGCACAAGTGC
AATCAGACCTGCCTGGACAGAATCGCCGCCGGCACCTTCAACGCCGGCGAGTTCAGCCTGCCCAC
CTTCGACAGCCTGAACATCACGGCCGCTAGCCTGAACGACGACGGCCTGGACAACCACACCATCCT
GCTGTACTACAGCACAGCCGCTAGCAGCCTGGCCGTGACCCTGATGCTGGCCATCTTCATCGTGTAC
ATGGTGAGCAGAGACAACGTGAGCTGCAGCATCTGCCTGtga

**Type/Subtype:** B/V

**Strain:** B/Austria/359417/2021
**Codon Optimization Scheme:** 1
**Source:** 3
ATGAAGGCCATCATCGTGCTGCTGATGGTGGTGACCAGCAACGCCGACCGCATCTGCACCGGCATC
ACCAGCAGCAACAGCCCCCACGTGGTGAAGACCGCCACCCAGGGCGAGGTGAACGTGACCGGCG
TGATCCCCCTGACCACCACCCCCACCAAGAGCCACTTCGCCAACCTGAAGGGCACCGAGACCCGC
GGCAAGCTGTGCCCCAAGTGCCTGAACTGCACCGACCTGGACGTGGCCCTGGGCCGCCCCAAGTG
CACCGGCAAGATCCCCAGCGCCCGCGTGAGCATCCTGCACGAGGTGCGCCCCGTGACCAGCGGC
TGCTTCCCCATCATGCACGACCGCACCAAGATCCGCCAGCTGCCCAACCTGCTGCGCGGCTACGAG
CACGTGCGCCTGAGCACCCACAACGTGATCAACACCGAGGACGCCCCCGGCGGCCCCTACGAGAT
CGGCACCAGCGGCAGCTGCCTGAACATCACCAACGGCAAGGGCTTCTTCGCCACCATGGCCTGGG
CCGTGCCCAAGAACAAGACCGCCACCAACCCCCTGACCATCGAGGTGCCCTACATCTGCACCGAG
GAGGAGGACCAGATCACCGTGTGGGGCTTCCACAGCGACGACGAGACCCAGATGGCCCGCCTGTA
CGGCGACAGCAAGCCCCAGAAGTTCACCAGCAGCGCCAACGGCGTGACCACCCACTACGTGAGCC

AGATCGGCGGCTTCCCCAACCAGACCGAGGACGGCGGCCTGCCCCAGAGCGGCCGCATCGTGGT
GGACTACATGGTGCAGAAGAGCGGCAAGACCGGCACCATCACCTACCAGCGCGGCATCCTGCTGC
CCCAGAAGGTGTGGTGCGCCAGCGGCAAGAGCAAGGTGATCAAGGGCAGCCTGCCCCTGATCGGC
GAGGCCGACTGCCTGCACGAGAAGTACGGCGGCCTGAACAAGAGCAAGCCCTACTACACCGGCGA
GCACGCCAAGGCCATCGGCAACTGCCCCATCTGGGTGAAGACCCCCCTGAAGCTGGCCAACGGCA
CCAAGTACCGCCCCCCCGCCAAGCTGCTGAAGGAGCGCGGCTTCTTCGGCGCCATCGCCGGCTTC
CTGGAGGGCGGCTGGGAGGGCATGATCGCCGGCTGGCACGGCTACACCAGCCACGGCGCCCACG
GCGTGGCCGTGGCCGCCGACCTGAAGAGCACCCAGGAGGCCATCAACAAGATCACCAAGAACCTG
AACAGCCTGAGCGAGCTGGAGGTGAAGAACCTGCAGCGCCTGAGCGGCGCCATGGACGAGCTGCA
CAACGAGATCCTGGAGCTGGACGAGAAGGTGGACGACCTGCGCGCCGACACCATCAGCAGCCAGA
TCGAGCTGGCCGTGCTGCTGAGCAACGAGGGCATCATCAACAGCGAGGACGAGCACCTGCTGGCC
CTGGAGCGCAAGCTGAAGAAGATGCTGGGCCCCAGCGCCGTGGAGATCGGCAACGGCTGCTTCGA
GACCAAGCACAAGTGCAACCAGACCTGCCTGGACCGCATCGCCGCCGGCACCTTCGACGCCGGC
GAGTTCAGCCTGCCCACCTTCGACAGCCTGAACATCACCGCCGCCAGCCTGAACGACGACGGCCT
GGACAACCACACCATCCTGCTGTACTACAGCACCGCCGCCAGCAGCCTGGCCGTGACCCTGATGAT
CGCCATCTTCGTGGTGTACATGGTGAGCCGCGACAACGTGAGCTGCAGCATCTGCCTGtga

**Strain:** B/Austria/359417/2021
**Codon Optimization Scheme:** 2
**Source:** 1, 2, 3
ATGAAGGCCATTATCGTGCTGCTGATGGTGGTGACCAGCAACGCCGACCGGATCTGTACCGGCATCA
CCAGCAGCAACAGCCCCCACGTGGTTAAGACCGCCACACAGGGCGAGGTGAACGTGACAGGCGTG
ATCCCTTTGACAACCACACCCACCAAGAGCCACTTCGCCAACCTGAAAGGCACCGAGACAAGAGGA
AAGCTGTGCCCCAAATGTCTGAATTGCACAGACCTGGACGTCGCCCTGGGCAGACCAAAGTGCACC
GGCAAGATCCCTAGCGCCAGAGTGTCCATCCTCCATGAAGTGCGGCCTGTAACCAGCGGCTGCTTC
CCCATCATGCACGACCGGACCAAGATTAGACAGCTGCCCAATCTGCTGAGAGGCTACGAGCATGTGA
GACTGAGCACCCACAACGTGATCAACACCGAGGATGCCCCCGGCGGTCCTTACGAGATCGGCACCA
GCGGCAGCTGTCTGAACATCACAAACGGAAAAGGCTTCTTCGCCACAATGGCTTGGGCCGTGCCTAA
GAACAAGACCGCTACCAACCCTCTGACAATCGAGGTGCCCTACATCTGCACCGAGGAAGAGGACCA
GATCACCGTGTGGGGCTTCCACAGCGATGATGAAACCCAAATGGCCAGACTGTACGGCGACAGCAA
GCCTCAGAAATTCACCTCCTCTGCTAATGGAGTGACCACCCACTACGTGTCCCAGATCGGCGGCTTC
CCTAATCAGACCGAGGACGGCGGACTGCCTCAAAGCGGCAGAATCGTGGTGGACTACATGGTGCAG
AAGTCCGGCAAAACAGGAACAATCACATACCAGAGGGGCATTCTGCTGCCACAAAAGGTGTGGTGCG
CCTCTGGCAAGTCTAAGGTGATCAAGGGCAGCCTGCCTCTGATCGGCGAAGCCGATTGCCTCCACGA
GAAGTACGGCGGCCTAAACAAGAGCAAGCCCTACTACACCGGCGAACACGCCAAGGCCATCGGAAA
CTGCCCTATCTGGGTGAAAACCCCTCTGAAGCTGGCCAACGGCACCAAGTATAGACCTCCAGCTAAG
CTGCTGAAGGAACGGGGCTTTTTCGGCGCCATCGCCGGTTTTCTGGAAGGCGGATGGGAGGGTATGA
TCGCCGGCTGGCACGGATATACAAGTCACGGCGCTCACGGCGTGGCCGTGGCTGCAGACCTGAAAA
GCACACAGGAGGCCATCAACAAAATCACCAAGAACCTGAATTCTCTGTCCGAGCTGGAAGTTAAGAAC
CTCCAGAGACTGAGCGGCGCCATGGACGAGCTGCACAATGAGATCCTTGAACTGGATGAAAAGGTCG
ACGACCTGCGGGCCGATACCATCAGCAGTCAGATCGAGCTGGCTGTGCTGCTGTCTAACGAGGGCAT
CATCAACAGCGAGGACGAGCACCTGCTGGCCCTGGAACGGAAGCTGAAAAAGATGCTGGGACCTAG
CGCCGTGGAAATCGGCAACGGCTGTTTCGAGACAAAGCACAAGTGCAACCAGACCTGCCTGGACAG
AATCGCCGCTGGCACATTTGATGCCGGAGAATTCAGCCTGCCTACCTTCGACAGCCTGAACATCACC
GCCGCCAGCCTGAACGACGATGGCCTGGACAACCACACCATCCTGCTGTACTACAGCACCGCCGCT
AGCAGCCTGGCCGTGACCCTGATGATCGCAATCTTCGTGGTCTACATGGTCTCTAGAGATAATGTGTC
CTGTTCAATTTGCCTGtga

**Strain:** B/Austria/359417/2021
**Codon Optimization Scheme:** 3
**Source:** 1, 3
ATGAAGGCCATCATCGTGCTGCTGATGGTGGTGACAAGCAACGCCGACAGAATCTGCACCGGCATCA
CAAGCAGCAACAGCCCCCACGTGGTGAAAACCGCGACACAAGGCGAGGTGAACGTGACCGGCGTG
ATCCCCCTGACCACCACCCCCACCAAGAGCCACTTCGCCAACCTGAAGGGCACCGAGACAAGAGG
CAAGCTGTGCCCCAAGTGCCTGAACTGCACCGACCTGGACGTGGCCCTGGGCAGACCCAAGTGCA
CCGGCAAGATCCCTAGCGCTAGAGTGAGCATCCTGCACGAGGTGAGACCCGTGACAAGCGGCTGCT
TCCCCATCATGCACGACAGAACCAAGATCAGACAGCTGCCCAACCTGCTGAGAGGCTACGAGCACG
TGAGACTGAGCACCCACAACGTGATCAACACCGAGGACGCCCCCGGCGGCCCCTACGAGATCGGC
ACAAGCGGCAGCTGTCTGAATATCACCAACGGAAAGGGCTTCTTCGCCACCATGGCCTGGGCCGTG
CCCAAGAACAAGACCGCTACGAACCCCCTGACCATCGAGGTGCCCTACATCTGCACCGAGGAGGAG
GATCAGATCACCGTGTGGGGCTTCCACAGCGACGACGAGACACAGATGGCTAGACTGTACGGCGAC
AGCAAGCCTCAGAAGTTCACAAGCAGCGCCAACGGCGTGACCACCCACTACGTGTCTCAGATCGGC
GGCTTCCCCAATCAGACCGAGGACGGCGGCCTGCCTCAGAGCGGCAGAATCGTGGTGGACTACATG
GTGCAGAAGAGCGGCAAGACCGGCACCATCACCTATCAGAGAGGCATCCTGCTGCCTCAGAAGGTG
TGGTGCGCTAGCGGCAAGAGCAAGGTGATCAAGGGCAGCCTGCCCCTGATCGGCGAGGCCGACTG
CCTGCACGAGAAGTACGGCGGCCTGAACAAGAGCAAGCCCTACTACACCGGCGAGCACGCCAAGG
CCATCGGCAACTGCCCCATCTGGGTGAAGACCCCCCTGAAGCTGGCCAACGGCACCAAGTACAGAC
CCCCCGCCAAGCTGCTGAAGGAGAGAGGATTCTTTGGAGCCATCGCCGGCTTCCTGGAGGGAGGCT
GGGAAGGTATGATAGCCGGGTGGCACGGCTATACTAGCCACGGCGCTCATGGCGTGGCCGTGGCC
GCCGACCTGAAGAGCACCCAAGAGGCCATCAACAAGATCACCAAGAACCTGAACAGCCTGAGCGAG
CTGGAGGTGAAGAACCTGCAGAGACTGAGCGGCGCCATGGACGAGCTGCACAACGAGATCCTGGAG
CTGGACGAGAAGGTGGACGACCTGAGAGCCGACACCATCAGCTCTCAGATCGAGCTGGCCGTGCTG
CTGAGCAACGAGGGCATCATCAACAGCGAGGACGAGCACCTGCTGGCCCTGGAGAGAAAGCTGAAG
AAGATGCTGGGCCCTAGCGCCGTGGAGATCGGCAACGGCTGCTTCGAGACCAAGCACAAGTGCAAT
CAGACCTGCCTGGACAGAATCGCCGCCGGCACCTTCGACGCCGGCGAGTTCAGCCTGCCCACCTT
CGACAGCCTGAACATCACCGCCGCGAGCCTGAACGACGACGGCCTGGACAACCACACCATCCTGC
TGTACTACAGCACCGCCGCTAGCTCCCTGGCCGTGACCCTGATGATCGCCATCTTCGTGGTGTACAT
GGTGAGCAGAGACAACGTGAGCTGCAGCATCTGCCTGtga

**Strain:** B/Washington/02/2019
**Codon Optimization Scheme:** 2
**Source:** 2
ATGAAGGCCATCATCGTGCTGCTGATGGTCGTGACAAGCAATGCCGACAGAATCTGTACCGGAATCAC
ATCCAGCAACAGCCCCCACGTGGTCAAGACCGCCACACAGGGCGAAGTCAACGTGACCGGCGTGA
TCCCCCTGACCACCACCCCTACAAAGAGCCACTTCGCCAATCTGAAGGGCACCGAGACAAGAGGCA
AGCTGTGCCCTAAGTGTCTGAATTGCACTGATCTGGACGTGGCCCTGGGCCGGCCAAAATGCACCGG
CAAGATCCCTTCTGCTAGAGTGTCCATCCTGCATGAGGTGCGGCCTGTGACCAGCGGCTGCTTCCCC
ATCATGCACGACCGCACCAAGATTAGACAGCTGCCTAACCTGCTTAGAGGCTACGAACATGTGCGGC
TGAGCACCCACAATGTGATCAACGCCGAGGACGCCCCTGGCAGGCCTTACGAGATCGGCACCAGC
GGCTCCTGCCCCAACATCACCAACGGAAATGGATTCTTTGCCACCATGGCTTGGGCCGTGCCCAAGA
ACAAGACCGCCACCAACCCTCTGACCATCGAGGTGCCTTACATCTGCACAGAAGGCGAGGATCAGAT
CACAGTGTGGGGCTTCCACTCTGATAGCGAGACACAGATGGCCAAGCTGTACGGCGACAGCAAGCC
GCAGAAATTCACCAGCTCTGCCAACGGCGTGACAACACACTACGTGTCGCAGATCGGCGGCTTCCCT
AACCAGACCGAGGACGGCGGACTGCCCCAAAGCGGCCGGATCGTCGTGGACTACATGGTGCAGAA

GTCTGGAAAGACCGGCACCATCACCTACCAGAGAGGAATTCTGCTGCCACAAAAAGTGTGGTGCGCC
AGCGGCAGATCTAAAGTGATCAAGGGCAGCCTCCCCCTGATCGGCGAAGCCGACTGTCTGCACGAA
AAGTACGGAGGCCTGAACAAGAGCAAGCCTTATTATACAGGCGAACACGCCAAGGCCATCGGCAACT
GCCCTATTTGGGTGAAGACCCCCCTGAAACTGGCCAACGGCACAAAATACAGACCACCTGCTAAGCT
CCTGAAGGAACGGGGATTTTTCGGCGCCATTGCCGGATTCCTGGAAGGCGGCTGGGAGGGCATGAT
CGCCGGCTGGCACGGATACACCAGCCACGGAGCCCACGGCGTGGCCGTAGCCGCAGACCTGAAG
TCCACCCAGGAGGCCATCAACAAGATCACCAAGAACTTGAACTCCCTGTCTGAACTGGAGGTGAAAAA
TCTCCAGCGGCTGTCTGGAGCTATGGACGAGCTGCACAACGAGATCCTGGAACTGGATGAGAAGGTG
GATGACCTGAGAGCCGATACAATCAGCTCTCAGATCGAGCTGGCCGTGCTGCTGAGCAACGAGGGC
ATCATCAACTCCGAGGATGAGCACCTGCTGGCCCTGGAAAGAAAGCTGAAAAAGATGCTGGGCCCTA
GCGCCGTGGAAATCGGCAATGGCTGTTTCGAGACAAAGCACAAGTGCAACCAGACCTGTCTGGACAG
AATCGCCGCTGGCACCTTCGATGCCGGCGAATTCAGCCTGCCTACCTTTGACAGCCTGAACATCACT
GCAGCTAGCCTGAACGACGACGGCCTGGACAACCACACCATCCTTCTGTACTACAGCACAGCTGCTT
CTAGCCTAGCTGTTACACTGATGATCGCCATCTTCGTGGTGTACATGGTGAGTAGAGATAACGTGAGCT
GCAGCATCTGCCTGtga
