## Supplemental Table 3 for "Single Platform Solution for Identity and Quantification of Influenza Hemagglutinin (HA) mRNA Constructs and Resulting Expressed Proteins for Application to Influenza mRNA Vaccines"

| | | H1 | | | | | | | | | H3 | | | | | | | | |
|---|---|---|---|---|---|---|---|---|---|---|---|---|---|---|---|---|---|---|---|
| | Strain | A/Wisconsin/67/2022 | | | | | | A/Sydney/5/2021 | A/Wisconsin/588/2019 | A/Michigan/45/2015 | A/Darwin/6/2021 | | | | | | A/Cambodia/e826360/2020 | A/HongKong/45/2019 | A/Singapore/INFIMH-16-0019/2016 |
| Vaccine Strain Year | 2018 SH | | | | | | | | | x | | | | | | | | | |
| | 2018-2019 NH | | | | | | | | | x | | | | | | | | | x |
| | 2019 SH | | | | | | | | | x | | | | | | | | | |
| | 2019-2020 NH | | | | | | | | | | | | | | | | | | |
| | 2020 SH | | | | | | | | | | | | | | | | | | |
| | 2020-2021 NH | | | | | | | | | | | | | | | | | x | |
| | 2021 SH | | | | | | | | x | | | | | | | | | x | |
| | 2021-2022 NH | | | | | | | | x | | | | | | | | x | | |
| | 2022 SH | | | | | | | | x | | x | x | x | x | x | x | | | |
| | 2022-2023 NH | | | | | | | | x | | x | x | x | x | x | x | | | |
| | 2023 SH | | | | | | | x | | | x | x | x | x | x | x | | | |
| | 2023-2024 NH | x | x | x | x | x | x | | | | x | x | x | x | x | x | | | |
| Codon optimization | | Method 3 | Method 3 | Method 3 | Method 2 | Method 2 | Method 1 | Method 2 | Method 2 | Method 4 | Method 2 | Method 2 | Method 2 | Method 3 | Method 3 | Method 1 | Method 2 | Method 2 | Method 4 |
| Manufacturer | | Source 1 | Source 2 | Source 3 | Source 2 | Source 3 | Source 3 | Source 2 | Source 2 | Source 1 | Source 1 | Source 2 | Source 3 | Source 1 | Source 3 | Source 3 | Source 2 | Source 2 | Source 1 |
| H1 Captures | H1-1 | 2.3 | 6.4 | 4.3 | 2.7 | 3.1 | 7.9 | 1.8 | 12.8 | 11.2 | 1.0 | 0.9 | 1.1 | 1.0 | 1.4 | 1.2 | 0.9 | 1.0 | 1.3 |
| | H1-2 | 7.6 | 16.8 | 8.1 | 3.0 | 5.6 | 16.7 | 1.2 | 1.1 | 1.5 | 1.0 | 0.9 | 1.0 | 1.0 | 1.2 | 1.1 | 0.9 | 0.9 | 1.0 |
| | H1-3 | 4.0 | 7.4 | 4.4 | 1.9 | 2.8 | 14.8 | 0.9 | 8.2 | 9.9 | 1.0 | 0.9 | 1.0 | 1.0 | 1.2 | 1.1 | 0.9 | 1.0 | 1.1 |
| | H1-4 | 1.1 | 1.1 | 1.1 | 1.5 | 2.3 | 1.0 | 16.5 | 1.0 | 1.0 | 1.0 | 1.0 | 1.0 | 1.0 | 1.1 | 0.9 | 0.9 | 1.0 | 1.0 |
| | H1-5 | 2.8 | 16.9 | 11.0 | 3.9 | 6.6 | 5.5 | 1.0 | 5.8 | 0.9 | 0.9 | 0.9 | 1.0 | 0.9 | 0.9 | 0.9 | 0.9 | 1.0 | 0.9 |
| | H1-6 | 15.8 | 16.8 | 13.0 | 1.3 | 1.6 | 17.0 | 1.9 | 16.2 | 1.6 | **3.5** | 2.3 | **3.1** | 1.1 | 1.7 | 1.6 | **3.3** | 1.4 | 1.0 |
| | H1-7 | 11.2 | 16.8 | 13.0 | 6.6 | 15.9 | 17.5 | 2.6 | 16.2 | 4.3 | 1.0 | 1.0 | 1.0 | 2.0 | **3.9** | 1.1 | 1.0 | 1.2 | 1.0 |
| | H1-8 | 3.1 | 5.9 | 7.4 | 1.2 | 1.8 | 4.6 | 7.3 | 1.4 | 1.2 | 1.0 | 1.0 | 1.0 | 0.9 | 1.1 | 0.9 | 0.9 | 1.1 | 1.0 |
| | H1-9 | 4.8 | 14.7 | 13.0 | 3.0 | 4.4 | 4.3 | 1.0 | 3.3 | 1.6 | 1.2 | 1.3 | 1.2 | 1.0 | 1.0 | 1.0 | 1.0 | 1.1 | 1.0 |
| | H1-10 | 7.1 | 16.8 | 13.0 | 1.0 | 1.0 | 17.5 | 8.6 | 1.1 | 1.6 | 1.0 | 0.9 | 1.1 | 2.3 | **15.2** | **4.0** | 1.0 | 1.4 | 1.0 |
| H3 Captures | H3-1 | 1.0 | 1.0 | 0.9 | 0.9 | 1.0 | 1.0 | 0.8 | 1.0 | 1.2 | 1.1 | 0.9 | 1.1 | 1.8 | 4.6 | 17.4 | 3.6 | 1.3 | 3.0 |
| | H3-2 | 0.9 | 0.9 | 0.9 | 0.8 | 0.9 | 1.0 | 0.9 | 0.9 | 1.0 | 1.0 | 1.1 | 1.3 | 0.8 | 1.1 | 0.9 | 17.7 | 17.5 | 1.0 |
| | H3-3 | 1.1 | 1.3 | 0.9 | 1.0 | 1.1 | 1.0 | 1.0 | 0.9 | 1.2 | 1.0 | 0.9 | 1.0 | 1.4 | 3.5 | 10.9 | 9.3 | 3.6 | 8.5 |
| | H3-4 | 1.0 | 1.3 | 0.8 | 0.9 | 1.1 | 1.0 | 0.8 | 1.0 | 1.3 | 1.2 | 2.3 | 5.3 | 2.6 | 8.3 | 17.4 | 0.9 | 2.5 | 3.9 |
| | H3-5 | 1.0 | 1.0 | 0.8 | 0.8 | 1.1 | 1.3 | 0.8 | 1.1 | 1.3 | 1.0 | 0.9 | 1.0 | 7.8 | 15.2 | 17.3 | 0.9 | 15.5 | 16.5 |
| | H3-6 | 1.1 | 0.9 | 1.1 | 0.9 | 1.0 | 1.0 | 0.9 | 0.9 | 1.1 | 5.6 | 6.2 | 12.6 | 0.9 | 1.2 | 0.9 | 0.9 | 1.0 | 1.0 |
| | H3-7 | 1.0 | 1.0 | 0.9 | 0.9 | 1.1 | 1.1 | 0.9 | 1.0 | 1.3 | 1.0 | 1.0 | 1.1 | 16.0 | 15.2 | 17.2 | 3.3 | 1.1 | 18.4 |
| | H3-8 | 0.9 | 0.9 | 0.9 | 0.8 | 1.0 | 1.0 | 0.9 | 0.8 | 1.0 | 0.9 | 0.9 | 0.9 | 4.2 | 14.7 | 10.6 | 0.9 | 7.2 | 5.1 |
| | H3-9 | 1.0 | 0.9 | 1.0 | 0.8 | 0.9 | 1.0 | 0.9 | 0.9 | 0.9 | 6.0 | 10.3 | 15.9 | 0.9 | 0.9 | 0.9 | 3.7 | 1.1 | 0.9 |
| | H3-10 | 1.3 | 2.3 | 1.3 | 0.8 | 0.9 | 1.3 | 1.7 | 0.9 | 1.3 | 5.8 | 4.7 | 5.3 | 11.0 | 15.2 | 17.3 | 10.2 | 10.7 | 18.5 |
| B/Y Captures | BY-1 | 1.0 | 1.0 | 1.0 | 1.0 | 1.1 | 1.0 | 0.9 | 0.9 | 1.3 | 0.9 | 0.9 | 1.0 | 1.0 | 1.1 | 1.1 | 1.0 | 1.0 | 1.0 |
| | BY-2 | 1.0 | 0.9 | 0.9 | 0.8 | 1.0 | 0.9 | 0.8 | 0.9 | 1.1 | 0.9 | 0.8 | 0.9 | 0.9 | 0.9 | 1.1 | 0.9 | 1.0 | 1.0 |
| | BY-3 | 0.9 | 0.9 | 0.9 | 0.8 | 1.0 | 0.9 | 0.8 | 0.9 | 1.1 | 1.0 | 1.0 | 0.9 | 2.1 | **9.7** | 2.0 | 1.0 | 1.0 | 1.0 |
| B/V Captures | BV-1 | 1.0 | 1.0 | 0.9 | 0.8 | 1.0 | 1.1 | 0.8 | 1.0 | 1.1 | 1.1 | 0.9 | 1.0 | 1.0 | 0.9 | **3.3** | 1.4 | 1.0 | 1.4 |
| | BV-2 | 0.8 | 0.9 | 0.9 | 0.8 | 1.0 | 0.9 | 0.9 | 0.9 | 1.0 | 1.0 | 0.8 | 1.0 | 0.8 | 1.1 | 0.9 | 0.9 | 1.0 | 0.8 |
| | BV-3 | 1.1 | 1.1 | **3.5** | 0.9 | 1.0 | 0.9 | 0.9 | 1.3 | 1.4 | 0.9 | 0.9 | 1.0 | 0.9 | 0.9 | 1.0 | 1.0 | 1.0 | 0.9 |
| Pan B Captures | Pan B-1 | 0.9 | 0.9 | 0.9 | 0.8 | 0.9 | 0.9 | 1.0 | 0.8 | 1.3 | 0.9 | 0.9 | 0.9 | 0.8 | 0.9 | 1.7 | 0.9 | 1.0 | 1.1 |
| | Pan B-2 | 0.9 | 0.9 | 0.9 | 0.9 | 0.9 | 0.9 | 1.0 | 0.8 | **4.6** | 0.9 | 0.9 | 0.9 | 0.9 | 1.0 | 1.8 | 0.9 | 1.1 | 1.4 |
| | Pan B-3 | 2.7 | **3.8** | **10.9** | 0.8 | 1.1 | 2.8 | 0.9 | 1.3 | 1.1 | 0.9 | 0.9 | 0.9 | 0.9 | 1.4 | 2.1 | 0.9 | 1.1 | 1.3 |
| | Pan B-4 | 1.0 | 1.6 | 1.1 | 0.8 | 0.9 | 2.0 | 0.9 | 0.9 | **14.8** | 0.9 | 0.9 | 0.9 | 0.8 | 1.2 | 1.0 | 0.8 | 1.0 | 0.9 |
| | Pan B-5 | 1.0 | 2.2 | 1.3 | 0.8 | 1.0 | 1.2 | 0.9 | 0.9 | 1.0 | 0.9 | 0.9 | 0.9 | 1.0 | 1.1 | 1.0 | 0.9 | 1.0 | 0.9 |
| | Pan B-6 | 0.9 | 1.0 | 1.3 | 0.8 | 1.0 | 1.0 | 1.0 | 1.2 | 1.6 | 0.9 | 0.8 | 1.2 | 0.8 | 1.0 | 1.1 | 0.9 | 1.1 | 1.7 |

| | | B/Yam | | | | | | B/Vic | | | | | | | |
|---|---|---|---|---|---|---|---|---|---|---|---|---|---|---|---|
| | Strain | B/Phuket/3073/2013 | | | | | | B/Austria/359417/2021 | | | | | | B/Washington/02/2019 | B/Colorado/06/2017 |
| Vaccine Strain Year | 2018 SH | x | x | x | x | x | x | | | | | | | | |
| | 2018-2019 NH | x | x | x | x | x | x | | | | | | | | x |
| | 2019 SH | x | x | x | x | x | x | | | | | | | | x |
| | 2019-2020 NH | x | x | x | x | x | x | | | | | | | | x |
| | 2020 SH | x | x | x | x | x | x | | | | | | | x | |
| | 2020-2021 NH | x | x | x | x | x | x | | | | | | | x | |
| | 2021 SH | x | x | x | x | x | x | | | | | | | x | |
| | 2021-2022 NH | x | x | x | x | x | x | | | | | | | x | |
| | 2022 SH | x | x | x | x | x | x | x | x | x | x | x | x | | |
| | 2022-2023 NH | x | x | x | x | x | x | x | x | x | x | x | x | | |
| | 2023 SH | x | x | x | x | x | x | x | x | x | x | x | x | | |
| | 2023-2024 NH | x | x | x | x | x | x | x | x | x | x | x | x | | |
| Codon optimization | | Method 3 | Method 3 | Method 3 | Method 4 | Method 1 | Method 4 | Method 2 | Method 2 | Method 2 | Method 3 | Method 3 | Method 1 | Method 2 | Method 4 |
| Manufacturer | | Source 1 | Source 2 | Source 3 | Source 1 | Source 3 | Source 3 | Source 1 | Source 2 | Source 3 | Source 1 | Source 3 | Source 3 | Source 2 | Source 1 |
| H1 Captures | H1-1 | 1.0 | 1.1 | 1.1 | 1.0 | 1.2 | 1.1 | 1.1 | 1.1 | 1.1 | 1.0 | 1.1 | 1.2 | 1.0 | 1.3 |
| | H1-2 | 1.0 | 1.0 | 1.1 | 1.0 | 1.1 | 1.0 | 1.0 | 1.0 | 1.0 | 1.0 | 1.0 | 1.0 | 0.9 | 1.2 |
| | H1-3 | 0.9 | 1.0 | 1.1 | 0.9 | 1.1 | 1.0 | 1.0 | 1.0 | 1.0 | 1.0 | 1.0 | 1.0 | 0.9 | 1.1 |
| | H1-4 | 0.9 | 1.0 | 0.9 | 1.0 | 0.9 | 1.0 | 1.0 | 0.9 | 1.0 | 1.1 | 1.0 | 0.9 | 1.0 | 1.0 |
| | H1-5 | 1.1 | 1.3 | 0.9 | 0.9 | 0.9 | 0.9 | 0.9 | 0.9 | 0.9 | 1.0 | 1.1 | 0.9 | 0.9 | 0.9 |
| | H1-6 | 1.8 | 2.6 | 1.9 | 1.0 | 1.4 | 1.0 | 1.0 | 1.0 | 1.1 | 1.1 | 1.3 | 1.1 | 1.0 | 1.2 |
| | H1-7 | 1.0 | 1.0 | 1.0 | 1.0 | 1.4 | 1.0 | 1.0 | 1.1 | 1.2 | 1.0 | 1.1 | 1.3 | 0.9 | 1.1 |
| | H1-8 | 0.9 | 1.0 | 0.9 | 0.9 | 0.9 | 0.9 | 0.9 | 0.9 | 0.9 | 1.0 | 0.9 | 0.9 | 0.9 | 0.9 |
| | H1-9 | 1.4 | 1.9 | 1.5 | 1.0 | 2.1 | 1.0 | 1.0 | 1.0 | 1.0 | 1.3 | 1.3 | 1.4 | 1.0 | 1.0 |
| | H1-10 | 1.0 | 1.4 | 2.9 | 1.0 | 1.7 | 1.2 | 1.0 | 1.0 | 3.0 | 1.0 | **17.1** | 1.9 | 1.0 | 1.0 |
| H3 Captures | H3-1 | 1.0 | 1.1 | 1.2 | 1.0 | 0.9 | 1.0 | 1.0 | 1.0 | 1.0 | 1.0 | 1.0 | 1.0 | 0.9 | 1.1 |
| | H3-2 | 0.9 | 1.0 | 0.9 | 0.9 | 0.9 | 0.9 | 0.9 | 0.9 | 0.9 | 1.0 | 0.9 | 0.9 | 0.9 | 0.9 |
| | H3-3 | 0.9 | 1.0 | 1.0 | 0.9 | 1.0 | 1.0 | 1.0 | 1.0 | 0.9 | 0.9 | 1.0 | 1.0 | 0.9 | 1.0 |
| | H3-4 | 0.9 | 0.9 | 1.0 | 1.0 | 1.3 | 1.0 | 1.0 | 0.9 | 0.9 | 0.9 | 1.3 | 1.3 | 0.9 | 1.0 |
| | H3-5 | 0.9 | 1.1 | 1.0 | 0.9 | 1.2 | 1.0 | 0.9 | 0.9 | 0.9 | 0.9 | 0.9 | 1.1 | 0.9 | 1.0 |
| | H3-6 | 0.9 | 1.2 | 1.1 | 1.0 | 0.9 | 0.9 | 1.0 | 0.9 | 1.0 | **28.0** | 1.0 | 0.9 | 0.9 | 1.0 |
| | H3-7 | 1.0 | 1.0 | 1.1 | 1.0 | 1.7 | 1.0 | 1.0 | 1.0 | 1.0 | 1.0 | 1.3 | 2.8 | 1.8 | 1.2 |
| | H3-8 | 0.9 | 1.0 | 0.9 | 0.9 | 0.9 | 0.9 | 1.0 | 0.9 | 0.9 | 0.9 | 0.9 | 0.9 | 0.9 | 0.9 |
| | H3-9 | 0.9 | 0.9 | 0.9 | 0.9 | 0.9 | 0.9 | 0.9 | 0.9 | 0.9 | 0.9 | 0.9 | 0.9 | 0.9 | 0.9 |
| | H3-10 | 1.0 | 0.9 | 0.9 | 0.9 | 1.2 | 1.0 | 0.9 | 0.9 | 0.9 | 0.9 | 1.0 | 0.9 | 0.9 | 1.0 |
| B/Y Captures | BY-1 | 0.9 | 1.0 | 1.0 | 13.3 | 6.1 | 4.3 | 1.0 | 0.9 | 1.0 | 0.9 | 1.0 | 1.0 | 1.0 | 1.0 |
| | BY-2 | 3.3 | 9.5 | 12.8 | 5.6 | 14.2 | 6.2 | 1.0 | 0.9 | 1.0 | 1.0 | 1.9 | 1.9 | 0.9 | 1.1 |
| | BY-3 | 1.7 | 2.4 | 2.5 | 1.6 | 5.0 | 1.3 | 1.6 | 1.6 | 2.1 | 1.1 | 1.3 | 1.9 | 0.9 | 1.8 |
| B/V Captures | BV-1 | 0.9 | 1.0 | 1.0 | 0.9 | 1.9 | 1.0 | 4.0 | 9.0 | 16.8 | 3.2 | 8.5 | 14.2 | 1.2 | 11.8 |
| | BV-2 | 0.9 | 1.2 | 0.9 | 0.9 | 0.9 | 1.1 | 3.1 | 4.4 | 12.4 | 7.3 | 20.0 | 18.1 | 19.5 | 3.4 |
| | BV-3 | 1.4 | 1.5 | 1.2 | 2.5 | 1.4 | 0.9 | 1.3 | 1.4 | 1.8 | 6.8 | 11.9 | 10.2 | 1.2 | 7.9 |
| Pan B Captures | Pan B-1 | 1.3 | 4.0 | 2.5 | 5.3 | 17.7 | 0.9 | 1.0 | 1.3 | 1.7 | 1.6 | 5.2 | 18.1 | 0.9 | 5.6 |
| | Pan B-2 | 2.5 | 10.2 | 10.2 | 3.4 | 17.0 | 1.7 | 1.0 | 1.4 | 2.2 | 3.0 | 16.0 | 10.7 | 1.0 | 4.9 |
| | Pan B-3 | 4.4 | 12.7 | 17.5 | 1.3 | 17.5 | 5.2 | 1.3 | 2.5 | 3.6 | 5.3 | 14.5 | 10.4 | 3.3 | 1.2 |
| | Pan B-4 | 8.4 | 13.7 | 19.5 | 19.2 | 17.7 | 1.0 | 3.6 | 18.9 | 18.4 | 14.2 | 20.0 | 12.1 | 18.2 | 9.2 |
| | Pan B-5 | 17.6 | 18.2 | 19.5 | 11.0 | 17.7 | 18.0 | 1.0 | 1.0 | 1.1 | 20.4 | 20.0 | 18.1 | 12.4 | 7.8 |
| | Pan B-6 | 5.7 | 18.2 | 12.8 | 12.4 | 6.4 | 1.0 | 1.3 | 1.4 | 1.5 | 6.9 | 11.7 | 3.5 | 1.0 | 3.6 |

**Supplementary Table 3. Summary of capture oligonucleotide reactivity and specificity across all mRNA constructs evaluated at 20 µg/mL.** Signal-to-background (S/B) ratios greater than 3 for intended targets are highlighted in grey, whereas off-target S/B ratios greater than 3 are indicated in bold.
