## Supplemental Table 4 for "Single Platform Solution for Identity and Quantification of Influenza Hemagglutinin (HA) mRNA Constructs and Resulting Expressed Proteins for Application to Influenza mRNA Vaccines"

**Supplementary Table 4. Precision and accuracy of selected capture oligonucleotides for LNP-encapsulated HA-encoding mRNA constructs.** Precision (%RSD) and accuracy (% expected values) of monovalent LNP-encapsulated mRNAs encoding A/Wisconsin/67/2022 (H1), A/Darwin/6/2021 (H3), B/Phuket/3073/2013 (B/Y), B/Austria/356417/2021 (B/V) (Method 1 codon optimized, Source 3). Values represent low, mid, and high concentrations across the quantification range. Values were determined from four technical replicates (n = 4).

| Capture Oligo | | Concentration (µg/mL) | Precision (% RSD) | Accuracy (% expected) |
|---|---|---|---|---|
| H1 Captures | H1-1 | 0.9 | 1.7 | 96.9 |
| | | 3 | 5.5 | 95.7 |
| | | 24 | 4.9 | 85.8 |
| | H1-2 | 0.5 | 0.4 | 97.1 |
| | | 6 | 5.3 | 90.6 |
| | | 24 | 7.9 | 83.4 |
| | H1-3 | 0.5 | 1.9 | 98.1 |
| | | 6 | 2.9 | 97.6 |
| | | 24 | 2.2 | 88.5 |
| | H1-5 | 2.4 | 2.1 | 92.6 |
| | | 12 | 1.6 | 98.1 |
| | | 24 | 1.5 | 90.3 |
| | H1-8 | 1.8 | 1.4 | 98.8 |
| | | 12 | 1.6 | 98.1 |
| | | 24 | 1.5 | 90.3 |
| H3 Captures | H3-1 | 0.2 | 1.4 | 93.6 |
| | | 6 | 8.3 | 92.9 |
| | | 24 | 7.5 | 88.3 |
| | H3-3 | 0.5 | 2.1 | 101.1 |
| | | 6 | 7.7 | 93.7 |
| | | 24 | 7.3 | 94.1 |
| | H3-5 | 0.1 | 1.1 | 92.3 |
| | | 9 | 7.3 | 92.1 |
| | | 24 | 6.1 | 90.7 |
| | H3-8 | 0.1 | 1.0 | 103.8 |
| | | 9 | 2.3 | 95.7 |
| | | 24 | 1.5 | 94.0 |
| B/Y Captures | BY-1 | 0.9 | 7.4 | 118.1 |
| | | 3 | 4.0 | 115.9 |
| | | 12 | 5.6 | 118.0 |
| | Pan B-4 | 0.5 | 3.8 | 107.4 |
| | | 3 | 4.1 | 104.5 |
| | | 12 | 4.2 | 106.6 |
| | Pan B-5 | 0.5 | 1.1 | 119.0 |
| | | 3 | 2.2 | 116.0 |
| | | 9 | 0.7 | 124.5 |
| | Pan B-6 | 0.9 | 1.5 | 117.3 |
| | | 12 | 2.3 | 119.0 |
| | | 24 | 3.1 | 119.2 |
| B/V Captures | BV-2 | 0.5 | 1.0 | 94.4 |
| | | 6 | 3.0 | 96.1 |
| | | 30 | 4.5 | 91.1 |
| | BV-3 | 0.5 | 3.1 | 100.6 |
| | | 6 | 3.7 | 96.4 |
| | | 24 | 0.4 | 93.5 |
| | Pan B-4 | 0.9 | 6.8 | 97.8 |
| | | 6 | 1.9 | 97.0 |
| | | 24 | 1.7 | 94.6 |
| | Pan B-5 | 0.06 | 3.1 | 95.7 |
| | | 6 | 2.1 | 102.1 |
| | | 24 | 1.9 | 89.3 |
| | Pan B-6 | 1.8 | 2.9 | 96.4 |
| | | 6 | 4.3 | 94.1 |
| | | 30 | 4.3 | 95.0 |
